# Sticking the landing: leg mechanosensory feedback coordinates the transition from flight to landing

**DOI:** 10.64898/2026.09.14.751322

**Authors:** Tse-Wei Kuo, Brooke Bingenheimer, Brooke Ferguson, Himadri Sunam, Francisca Bermedo-Garcia, Sweta Agrawal

**Affiliations:** School of Neuroscience, Virginia Tech, Blacksburg, VA, USA; Department of Zoology, The University of British Columbia, Vancouver, British Columbia, Canada

**Keywords:** sensorimotor integration, flight, landing, behavioral coordination, *Drosophila melanogaster*, mechanosensory feedback

## Abstract

Landing is an especially demanding stage of flight. Flying animals must rapidly transform multimodal sensory information into the coordinated movement of multiple limbs in order to transition from flight to a stable standing posture, often on surfaces that vary in stability, texture, and orientation. Failure to do so can be costly, leading to injury or death. Here, we examine the role of mechanosensory feedback in guiding landing in the fruit fly, *Drosophila melanogaster*. We describe a novel behavioral assay in which we deliver localized mechanical contact to individual legs of tethered, flying flies while simultaneously quantifying their three-dimensional behavioral kinematics. We find that mechanical contact of the distal legs reliably induces flight cessation, and initial leg contact triggers distinct patterns of inter-leg coordination. Recruitment of subsequent leg contacts predicts landing likelihood, suggesting that coordinated feedback from multiple legs reinforces the landing transition. Next, we combine synapse-resolution connectomes with optogenetic manipulations to identify the mechanosensory pathways underlying these behaviors. Finally, we describe a population of inhibitory ascending neurons that integrates mechanosensory input from the legs and facilitates the transition from visually-mediated landing initiation to touch-mediated flight cessation. Overall, our results reveal an important set of neural circuits that convert local tactile feedback into a coordinated, rapid behavioral state transition.

## Introduction

Flying animals regularly transition between terrestrial and aerial environments to accomplish survival behaviors like feeding, mating, or resting^1–3^. Uncontrolled or failed landings during these transitions can be costly, resulting in an excessive expenditure of energy, injury, or death. As a result, performing a successful landing is arguably the most important step of flight.

Work across species has revealed that landing is a complex behavior that occurs via a well-coordinated sequence of sub-behaviors, including body-deceleration, body rotations, leg extension, grasping, and flight cessation (**Figure 1A**)^2,4–7^. Much of this previous work has focused on how visual feedback coordinates landing behaviors so that animals are in the correct position and posture at the correct time to facilitate a controlled landing. Less attention has been paid to the actual moment of touch down and the subsequent transition to standing. During most landings, the legs are the first point of contact with the landing substrate^2,8^. The animal must then then coordinate across its multiple limbs to adhere to the surface and transition to a stable standing posture. This final step is crucial for a successful landing, can happen rapidly (within 50 ms in the case of some insect species^9^), and is typically quite robust, with many species able to land on substrates that vary considerably in their stability, texture, and orientation^10^. Thus, landing is not only an important flight behavior, but also a useful model to understand how sensorimotor circuits rapidly and precisely coordinate movements across multiple limbs to effect behavioral transitions.

**Figure 1.**
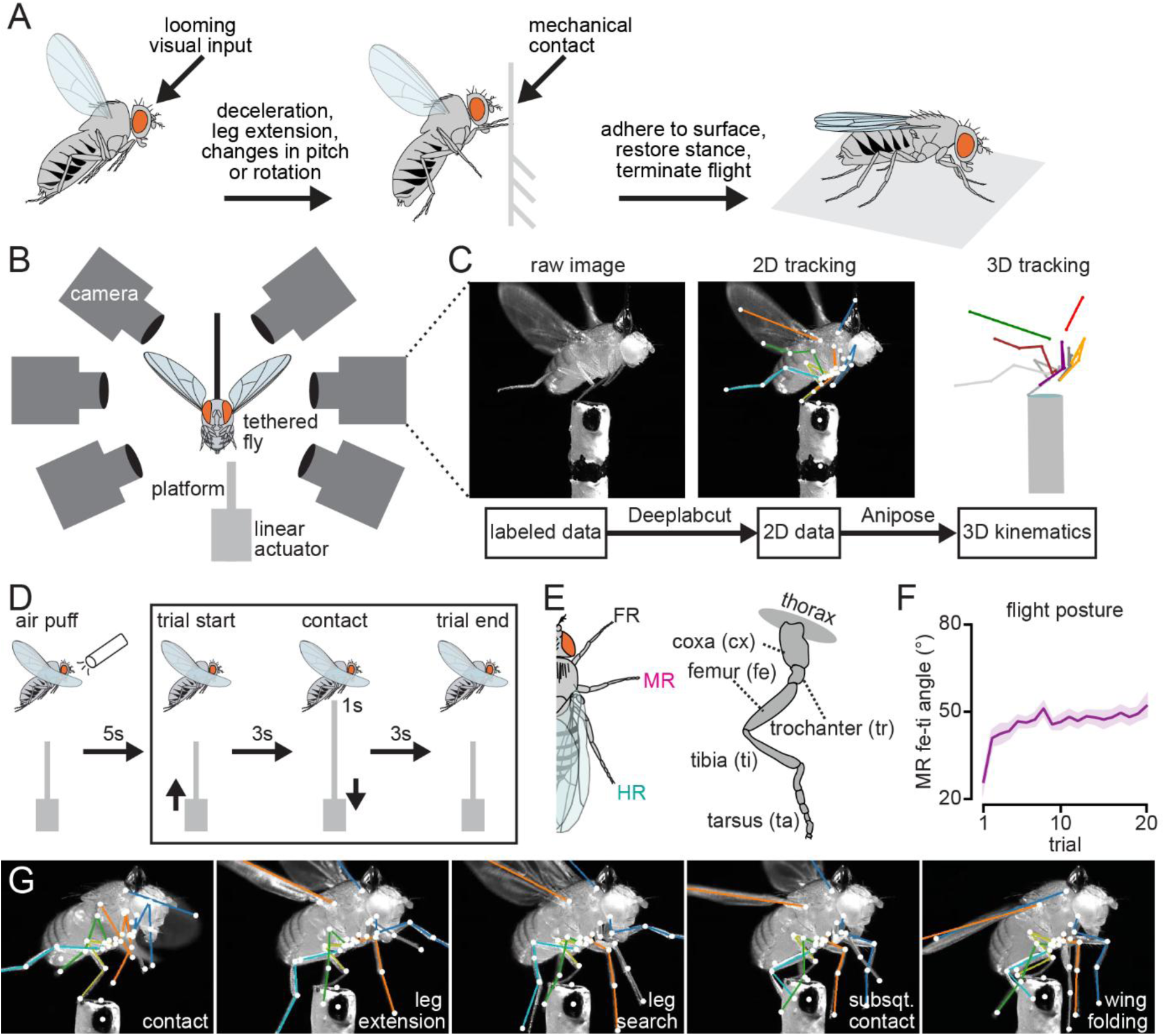
A novel behavioral assay to study contact-induced landing transitions. **(A)** Landing consists of a sequence of sub-behaviors, including body-deceleration, body rotations and/or pitching, leg extension, grasping, and flight cessation. Different sensory cues are thought to trigger different sub-behaviors. **(B)** Schematic of the behavioral assay. Six synchronized cameras recorded the behavior of tethered flies while a motorized platform delivered mechanical stimulation from below. **(C)** We trained a DeepLabCut^38^ network to estimate the fly’s 2D pose in each of our videos. Multi-view tracking was then reconstructed into 3D kinematics using Anipose^39^. **(D)** Timeline of a typical behavioral trial. A gentle air puff was delivered 5 s before trial onset to initiate flight. Over the first 3 s of each trial, the motorized platform extended to reach the target position, remained stationary for 1 s, and then retracted over the final 3 s. **(E)** *Drosophila* leg anatomy. Left: front right (FR), middle right (MR), and hind right (HR) legs are shown. Right: The different leg segments include the coxa (cx), trochanter (tr), femur (fe), tibia (ti), and tarsus (ta). **(F)** Mean (dark line) and standard error (shaded lines) femur-tibia joint angle of the contacted middle right leg across 20 consecutive trials in the 1s preceding platform contact (N=14 flies). **(G)** Representative images of some of the common behaviors we observed after platform contact.

Mechanosensory neurons on the legs have been shown to encode surface contact and the distribution of body load^11^. These neurons are therefore likely key to coordinating successful landings. However, while the role of these mechanosensory neurons has been well-described for behaviors like walking, their function during landing is still largely unknown. Here, we developed a novel behavioral assay using the fruit fly, *Drosophila melanogaster*, to determine how feedback from leg mechanosensory neurons coordinates limb movements during landing. We induced landing in tethered, flying flies by elevating a platform to contact their legs, mimicking the contact that occurs during landing, while simultaneously tracking the 3D kinematics of leg joints, wings, and body posture of flies as they responded to this contact. By adjusting the location of the contact, we dissected how mechanical perturbation of the legs alters limb coordination and flight cessation. We then used connectivity analyses from a synapse-resolution volume of the fly central nervous system (CNS) to guide optogenetic experiments manipulating subsets of leg mechanosensory neurons. Finally, we identify an ascending neuron that coordinates the transition from flight to landing. Altogether, these experiments provide novel insights into how mechanosensory feedback rapidly coordinates ethologically important behaviors, enabling flying animals to smoothly transition between locomotor modes.

## Results

### A novel behavioral assay to manipulate mechanosensory feedback in landing

We delivered mechanical stimulation to individual legs of tethered, flying flies via a motorized platform (**Figure 1B-D**). All experiments were performed in the dark with infrared (IR) illumination so as to eliminate visual cues, and we recorded the flies’ behavior via six high-speed cameras. Using these videos, we reconstructed the three-dimensional kinematics of the legs, wings, and body and aligned them to the platform’s movement (**Figure 1C**). Each fly experienced 20 contact events, or trials, with a 10 s intertrial interval. Between each trial, an air puff aimed at the right anterior side of the fly was provided to ensure the fly was flying at trial start (**Figure 1D**). Across repeated trials, we observed a consistent shift in each fly’s flight posture prior to platform contact characterized by a gradual increase in the femur-tibia (fe-ti) joint angle (**Figure 1E-F**). This shift suggests that flies adapt to repeat mechanical stimulation by slightly extending their legs after the initial few trials. To account for this adaptation, and because we worried that changes in leg posture would reduce the stereotypy of our mechanical stimulus, we excluded the first three trials from all subsequent analyses. Following mechanical contact by the platform, flies exhibited a consistent array of behaviors, including leg extension, rhythmic searching leg movements, subsequent contacts of the platform by the non-contacted legs, flight cessation, and folding of the wings (**Figure 1G**). We defined landing as flight cessation, including the folding of wings, that happened within 0.71 s after contact by the platform (See Methods for more details on how this threshold was selected).

### Mechanosensory contact of the legs causes flies to stop flying

We first tested how the location of the platform contact affected the fly’s behavior. Flies rarely stopped flying if the platform was raised without contacting the fly (**Figure 2A**). Contacting the tibia-tarsus (ti-ta) joint caused flies to stop flying during the majority of trials whereas contact of the coxa-trochanter (cx-tr) joint led to significantly fewer landings (**Figure 2A, Movie S1**). In addition, the landings that did occur upon contact of the cx-tr joint happened with a greater latency compared to those after contacting the ti-ta joint (**Figure 2B**). We also found that landing responses were largely consistent regardless of which leg (front, middle, or hind) was contacted (**Figure 2A**), even though mechanical contact led to different patterns of joint angle changes of each of these legs (**Figure 2C**). This variability in leg movement occurs because the three pairs of legs are held in distinct postures during flight and therefore differently moved by a contact made to the same joint. Contacting the fly’s abdomen also led to minimal landing response similar to contacting the cx-tr leg joint (**Figure 2A**). Moreover, we noticed that during these trials, even though the primary point of contact was the abdomen, the hind legs would also frequently contact the platform. When we removed the hind legs, contacting the abdomen no longer caused the flies to stop flying (**Figure 2A**). Overall, these results reveal that contact location influences both whether landing occurs and the speed with which the landing transition is executed.

**Figure 2.**
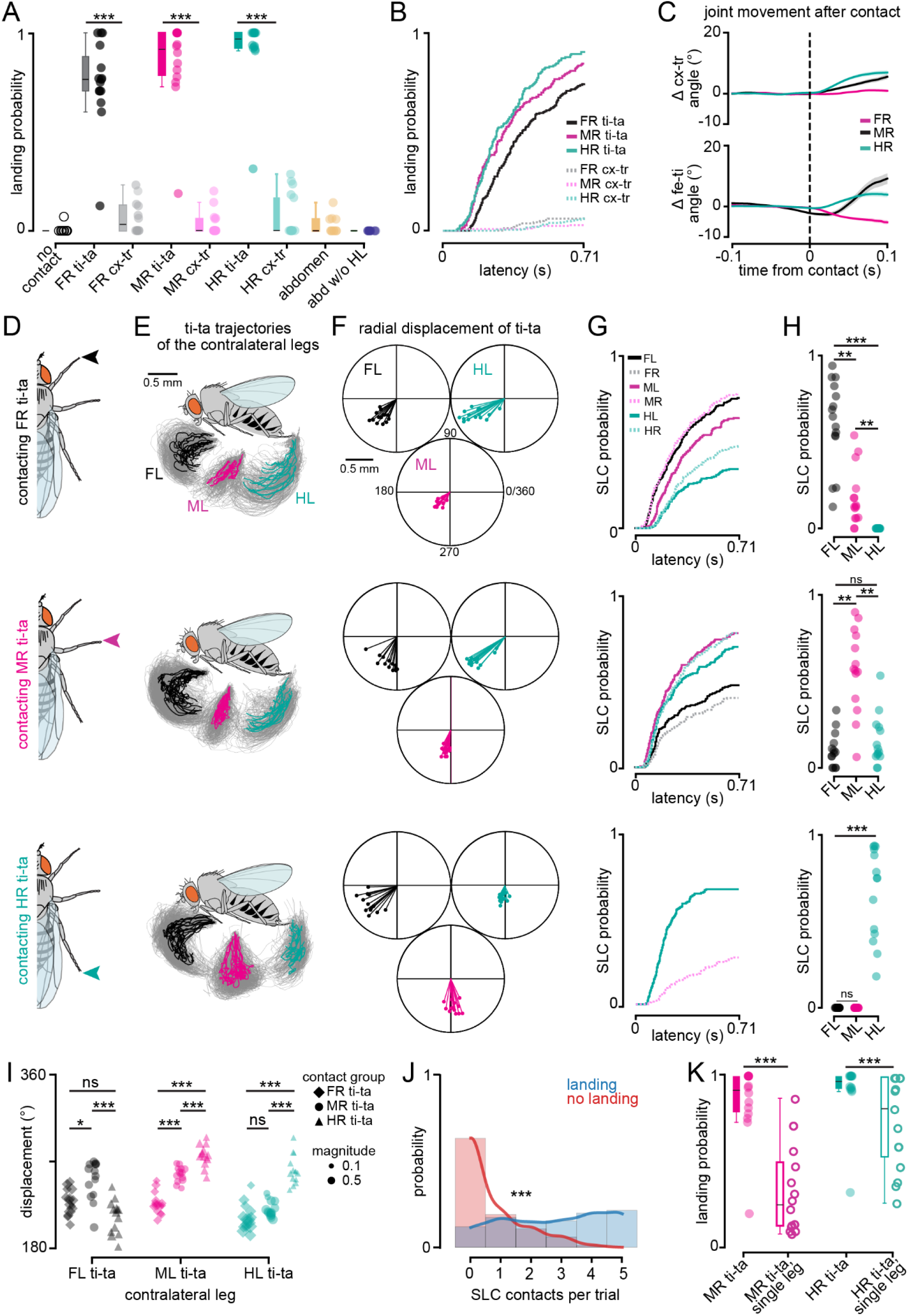
Mechanical contact of the legs leads to landing and stereotyped movements of the other legs. **(A)** Fly-wise landing probability following mechanical stimulation of the cx-tr and ti-ta joints of the front, middle, and hind right legs. Control groups include raising the platform without contacting the fly (no contact), contacting the fly’s abdomen, and abdomen contact with the hind legs removed (abd w/o HL). Circles represent the total fraction of trials per fly in which the fly was classified as landing. Statistical comparisons were performed using two-sided unpaired permutation tests on fly-wise landing probability. Box plots show the median (center line), interquartile range (box), and the most extreme values within 1.5 times the interquartile range (whiskers). Sample sizes, from left to right, are N=10, 15, 12, 14, 17, 14, 17, 11, and 10 flies, \*\*\**p*<0.001. **(B)** Cumulative landing probability following cx-tr (dashed lines) and ti-ta (solid lines) stimulation of the FR (gray), MR (magenta), and HR (teal) legs. Statistical comparisons were performed using two-sided unpaired permutation tests on fly-wise restricted mean survival time (RMST). Sample sizes same as (A). **(C)** Mean (solid line) and standard error of mean (shaded lines) of joint angle changes of the contacted leg following ti-ta stimulation. Top: change in cx-tr angle, bottom: change in fe-ti angle. FR (black): n=226 trials (top), 227 trials (bottom), MR (magenta): n=190 trials (top), 184 trials (bottom), and HR (teal): n=191 trials (top), 154 trials (bottom). **(D)** Schematic of contacted leg associated with each row’s data. **(E)** Contralateral ti-ta joint trajectories following stimulation of the leg indicated in (D). Individual trajectories are plotted in gray, and a per-fly average is shown in black (front leg, N=15 flies), pink (middle leg, N=14 flies) or teal (hind leg, N=14 flies). **(F)** Radial displacement of the contralateral ti-ta joint trajectories. Statistical comparisons were performed using two-sided fly-wise two-dimensional displacement-vector permutation tests. **(G)** Cumulative probability of subsequent leg contact (SLC) for each non-contacted leg following the initial platform contact. From top to bottom, N=15, 14, 14 flies and n=233, 190, 191 trials. Statistical comparisons were performed using two-sided unpaired permutation tests on fly-wise RMST. **(H)** Fly-wise probability that each contralateral leg was the first leg to establish a valid SLC. Statistical comparisons were performed using two-sided unpaired permutation tests on fly-wise first SLC probability. For FR, MR, and HR-ti-ta stimulation groups, N=15, 14, and 14 flies, respectively. ns: not significant, *p*≥0.05; *p<0.05; \*\**p*<0.01; \*\*\**p*<0.001. **(I)** Fly-wise mean projected ti-ta endpoint displacement angles are shown for each left ti-ta joint. Each point represents one fly. Point color indicates the leg joint (black: FL ti-ta, magenta: ML ti-ta, teal: HL ti-ta). Marker shape indicates the contralateral leg that was contacted (diamond: FR ti-ta, circle: MR ti-ta, triangle: HR ti-ta). Marker size represents the magnitude of the fly-mean displacement vector. Angles were calculated from the fly-mean two-dimensional displacement vector and plotted in degrees. Directional differences between contact groups were assessed using two-sided fly-wise directional displacement permutation tests. **(J)** Distribution of the number of SLC events in landing and no landing trials, Blue line/bars indicate landing trials (n = 509), and red line/bars indicate no landing trials (n = 88); bar height represents the probability of trials with the indicated number of SLC events. SLC counts were compared between landing and no landing trials using two sided unpaired permutation test at the trial level, \*\*\**p*<0.001. **(K)** Fly-wise landing probability of intact flies and flies with non-contacted legs surgically ablated. Sample sizes, from left to right: N=15, 14, 17, and 14 flies. Unpaired permutation tests were performed on the difference in fly-wise mean landing probability. \*\*\**p*<0.001. Exact *p* values and additional statistical comparisons for Figure 2 are provided in Supplementary Table S1.

We noticed that upon contact of one leg, flies frequently make searching maneuvers with their other legs and move them to also contact the platform. These leg movements changed in their direction depending on the location of the contact (**Figure 2D-E**). To quantify these differences, we compared the displacement vectors of each of the legs on the side contralateral to the platform contact. We found that the direction of post-contact left leg movements differed significantly depending on which leg was contacted on the right side (**Figure 2F, I**). When we quantified the timing and identity of subsequent leg contacts (SLCs) made onto the platform following the initial mechanical stimulus, we found that distinct coordination patterns emerged across stimulation groups (**Figure 2G-H**). Contact of the front leg ti-ta joint led to SLCs by primarily the contralateral front and ipsilateral middle legs. In contrast, contact of the hind leg ti-ta joint led to engagement of the contralateral hind leg and ipsilateral middle leg. Together, these results indicate that the location of the initial mechanical stimulus reorganizes how the remaining legs move toward the platform.

The contact-dependent differences in SLCs raised the possibility that mechanical contact by multiple, coordinated legs could reinforce landing such that more SLCs lead to a higher likelihood of landing. In support of this possibility, we found that trials that resulted in successful landing were associated with more SLCs (**Figure 2J**). When we tested flies in which we ablated all legs except for the leg that we mechanically stimulated, thereby removing the possibility of SLCs, we found that landing probability was reduced but not completely abolished (**Figure 2K**). Thus, mechanical input from a single leg is sufficient to evoke flight cessation, but the availability of additional leg contacts substantially enhances landing performance.

Together, these results demonstrate that initial contact location influences both landing outcome and the pattern of subsequent inter-leg coordination. Furthermore, subsequent contact by additional legs is associated with increases in landing probability. These findings support a model in which successful landing depends not solely on the initial mechanical stimulus, but also on the coordinated recruitment of additional leg-mediated sensory feedback.

### Variability in leg contact impacts landing coordination and success

We noticed that substantial variability in landing success occurred even within the same stimulation condition contacting the same joint of the same leg, suggesting that factors beyond contact location influence the animals’ landing performance. When we examined trials in which we contacted the ti-ta joint of the front right leg, we found that contacting the joint sometimes led to an initial transient touch (TT) (**Figure 3A**) that resulted in distinct post-contact movements of the contacted leg’s fe-ti joint (**Figure 3B**). TT trials were less likely to lead to successful landing and were associated with longer landing latencies (**Figure 3C-D**).

**Figure 3.**
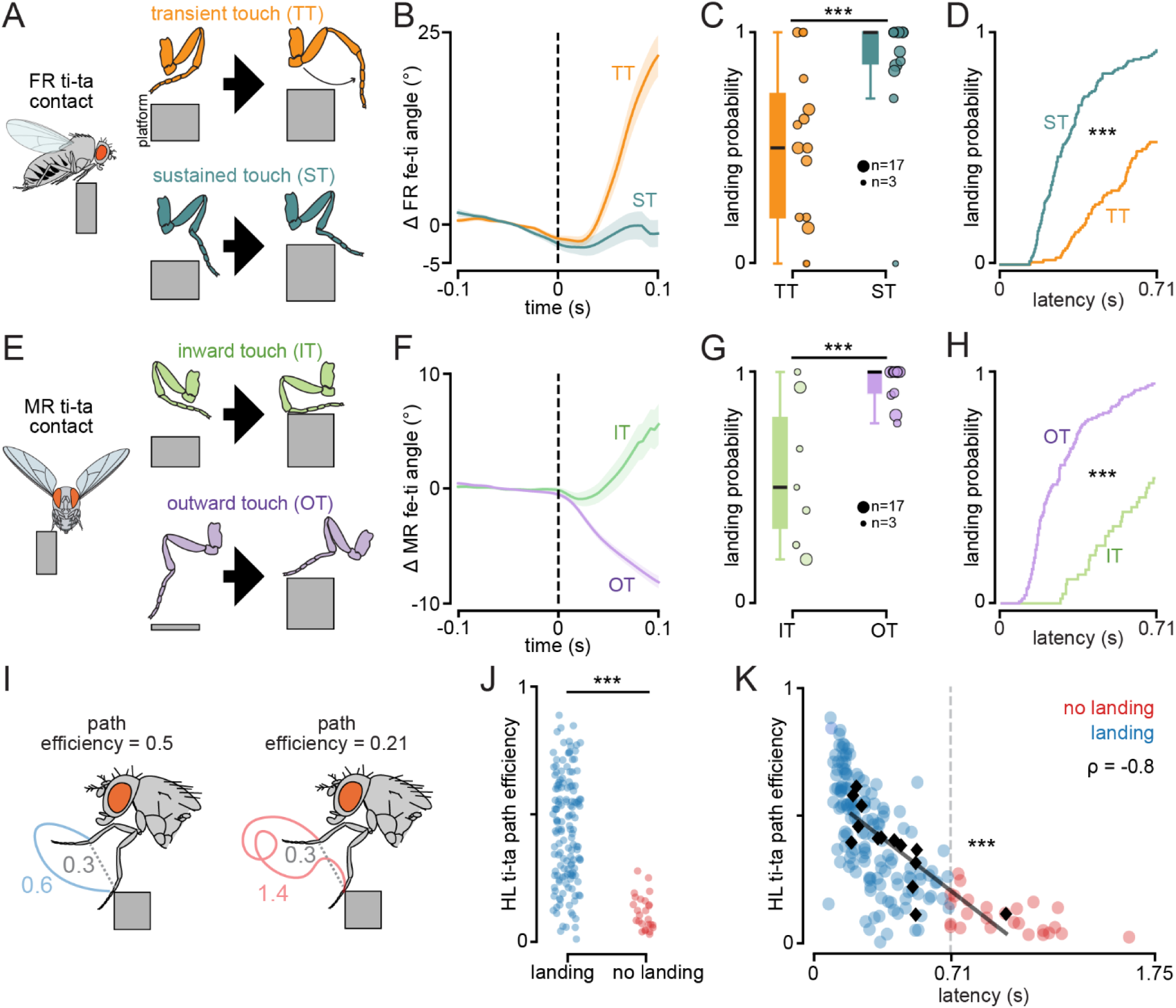
Subtle differences in platform contact contribute to landing variability. **(A, E)** Schematics illustrating different platform contacts made during ti-ta stimulation of the front (A) or middle (E) right legs. **(B, F)** The change in fe-ti joint angle of the contacted leg aligned to the moment of contact. Solid lines and shaded regions represent the mean ± SEM across trials. **(B)** Transient touch trials (TT, orange, n=100) are characterized by an increase in fe-ti joint angle compared to sustained touch trials (ST, dark blue, n=127). **(F)** Inward touch (IT, green, n=42) trials are characterized by a small increase in fe-ti joint angle whereas outward touch (OT, purple, n=142) trials are characterized by a decrease in fe-ti joint angle. **(C, G)** Fly-wise landing probability for each contact geometry. Dot size indicates the number of contributing trials for each fly. Differences in landing probability between contact geometries were assessed using two-sided trial-level label-shuffle permutation tests on binary landing outcomes, \*\*\**p*=5×10^-5^. **(D, H)** Cumulative probability of landing following moment of contact for each contact geometry. TT: n=101, ST: n=132, IT: n=48, OT: n=142. Landing-latency distributions were compared using two-sided log-rank tests, \*\*\**p*=2.3×10^-14^. **(I)** Schematic illustrating calculation of ti-ta trajectory path efficiency. **(J)** Path efficiency of the HL ti-ta joint trajectory in landing (blue, n=146) and no landing (red, n=26) trials from trials in which the MR ti-ta was contacted. Statistical comparisons were performed using two-sided unpaired permutation tests on trial-level path-efficiency values, \*\*\**p*=5×10^-5^. **(K)** HL ti-ta joint trajectory path efficiency relative to landing latency for trials in which the MR ti-ta was contacted. n=172 trials, N=14 flies, individual trials were assessed using Spearman’s rank correlation. ρ indicates the strength and direction of monotonic relationship between path efficiency and latency, \*\*\**p*=9.3×10^-39^.

We found similar variability when examining trials in which we contacted the ti-ta joint of the middle right leg. Even when contacting the same ti-ta joint of the same middle leg in the same fly, mechanical contact could cause the tibia to bend either towards the coxa (inward touch, IT) or away from the coxa (outward touch, OT) (**Figure 3E**), resulting in distinct post-contact femur-tibia (fe-ti) joint trajectories that represent mechanically distinct substrate interactions (**Figure 3F**). We found that, once again, these differences in contact geometry were associated with differences in landing performance and landing latency (**Figure 3G-H**).

Finally, we found that during trials in which the fly did not land within the predetermined threshold, the fly’s behavior suggested that it still perceived the mechanical contact because its other legs would make rhythmic, searching maneuvers. However, this searching behavior rarely resulted in legs successfully contacting the platform, as quantified by a lower path efficiency score (**Figure 3I-J**). Leg path efficiency is negatively correlated with landing latency at the trial level, demonstrating that more efficient movement organization is associated with faster landing transitions (**Figure 3K**).

Overall, these findings indicate that variability in landing performance is associated with differences in contact geometry, post-contact leg recruitment, and movement organization. Successful landing trials were characterized by greater subsequent leg engagement and more efficient leg movement trajectories.

### Connectomics predict subtype-specific differences in mechanosensory feedback during landing

Our behavioral experiments revealed not only that mechanosensory contact of the legs is sufficient and possibly required to cause flight cessation, but also that the location of this contact and how it moves the leg is important as well. These results suggest that activation of only specific subpopulations of leg mechanosensory neurons is likely to lead to landing and flight cessation. These neurons may be sensitive to mechanical contact (i.e., exteroceptive), changes in leg posture (i.e., proprioceptive), or both.

The fly leg is equipped with multiple mechanoreceptor types, including tactile bristles, chordotonal organ neurons, hair plate neurons, and campaniform sensilla^11^. Each of these mechanoreceptors is sensitive to a particular range of stimuli, though the same stimulus can also co-activate multiple receptor types. In an effort to identify mechanosensory neurons important for landing and flight cessation, we utilized FANC, a connectome generated from an electron microscopy-imaged volume of a female *Drosophila* ventral nerve cord (VNC)^12^. The VNC is analogous to the vertebrate spinal cord and includes axons from sensory neurons distributed throughout the body, motor neurons controlling movement of the wings, legs, and body, interneurons, and ascending and descending neurons connecting the VNC with the brain. We reasoned that mechanosensory neurons involved in landing and flight cessation should demonstrate downstream direct or indirect connectivity with flight premotor circuits. We identified axons associated with each mechanoreceptor type from the fly’s left front leg (**Figure 4A-D**), and examined their connectivity with wing motor neurons sorted into functional motor modules as identified by Lesser et al., (2024)^13^ (**Figure 4E**). Given the rapid nature of the flight cessation response, we focused on monosynaptic and disynaptic connections. We found that leg chordotonal organ neurons have no direct or indirect connectivity with wing motor circuits, and hair plate neurons are only very weakly connected to wing motor circuits. In contrast, tactile bristles and campaniform sensilla (CS) demonstrate much greater connectivity with wing motor circuits (**Figure 4F**). Most of this connectivity is indirect via a layer of interneurons, though a few leg CS neurons do synapse directly onto wing steering motor neurons. These direct connections are made by bilaterally-projecting CS neurons present on the trochanter^14^ onto wing steering motor neurons. Thus, connectivity analyses suggest that leg tactile bristles and CS likely mediate the flight cessation response seen during landing.

**Figure 4.**
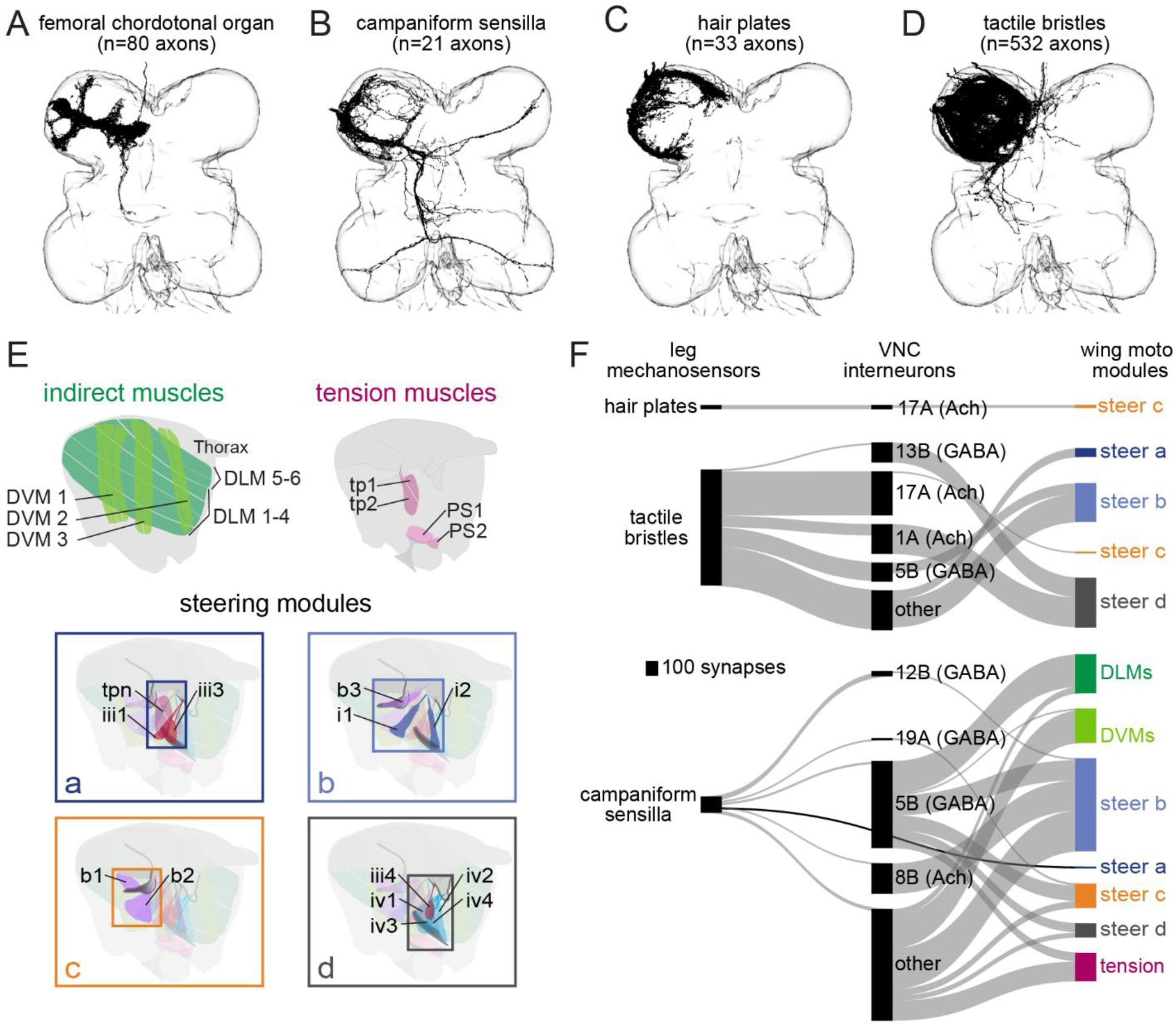
Connectivity analyses suggest that tactile bristles and campaniform sensilla mediate contact-induced landing. **(A-D)** Reconstructed mechanosensory axons from each subtype in the front left leg neuromere of the FANC connectomes (from left to right, n=80, 21, 33, 532 neurons). **(E)** Schematic of the different wing motor modules as classified by Lesser et al., (2024)^13^, figure modified from original with permission. These include the indirect muscles, tension muscles, and 4 subgroups of steering muscles. **(F)** Sankey plots demonstrating mono-and di-synaptic connectivity between leg mechanosensory neurons (left) and wing motor neurons (right). Direct connections are plotted in black, indirect connections in gray. VNC interneurons are grouped via their hemilineage, and their putative neurotransmitter is indicated based on Lacin et al., (2019)^44^. Neurons whose hemilineage could not be confidently identified are categorized as “other.” Only premotor and motor neurons that received at least 4 synapses from leg mechanosensory axons and only premotor neurons that supply at least 4 synapses to motor neurons are displayed.

### Optogenetic experiments confirm the role of bristles and CS during landing

We next tested whether activation of subsets of leg mechanosensory neurons contributes to contact-mediated landing as predicted by their synaptic connectivity to wing motor neurons. We genetically expressed the optogenetic channel csChrimson^15^ via driver lines that labeled different subsets of mechanosensory neurons, and then measured whether activation of these sensory neurons was sufficient to cause flight cessation. These genetic driver lines included iav-GAL4 which labels chordotonal organ neurons throughout the body^16^, a split-GAL4 line that labels hair plate neurons specifically from the leg^17^, eyg-GAL4 which labels CS throughout the body^18,19^, and 38b08-LexA combined with an intersectional genetic strategy developed by Elabbady et al. (2026) to label tactile bristles on the leg tarsus^20^. We also used a collection of split-GAL4 driver lines to target groups of CS neurons specifically on the leg. Across almost all genotypes tested, including wildtype (*canton-s*) flies, we consistently observed a transient leg extension response following light offset, suggesting a stereotyped behavioral response to termination of the red light stimulus independent of the specific neurons targeted. This response occurred after light offset and therefore did not affect our analysis of behavior during the trial.

Activation of chordotonal organ neurons or hair plate neurons did not significantly change flight cessation probability relative to controls (**Figure 5A, Movie S2**). In contrast, activation of tarsal tactile bristles or all CS increased flight cessation likelihood. Because the genetic driver line that broadly labels all leg CS (eyg-GAL4) also labels CS on the wings, halteres, and antennae, we next tested a collection of split-GAL4 lines that label smaller subsets of CS primarily on the legs. Of these, activation of the bilaterally-projecting leg CS via three different lines or tarsal CS did not lead to flight cessation. This result surprised us, because the bilateral CS axons reconstructed in FANC make direct, though weak, synapses onto wing motor neurons (**Figure 4F**). Activation of a line labeling more proximally-located CS present on the femur and trochanter of the leg did lead to an increased probability of flight cessation, but only one of two such lines tested led to a significant increase in flight cessation, and only when laser power was increased to 0.121 mWmm^-2^ (**Figure 5A**). Interestingly, activation of the tarsal bristles not only led to flight cessation in almost all trials, but the onset of this response was very rapid as well (**Figure 5B**).

**Figure 5.**
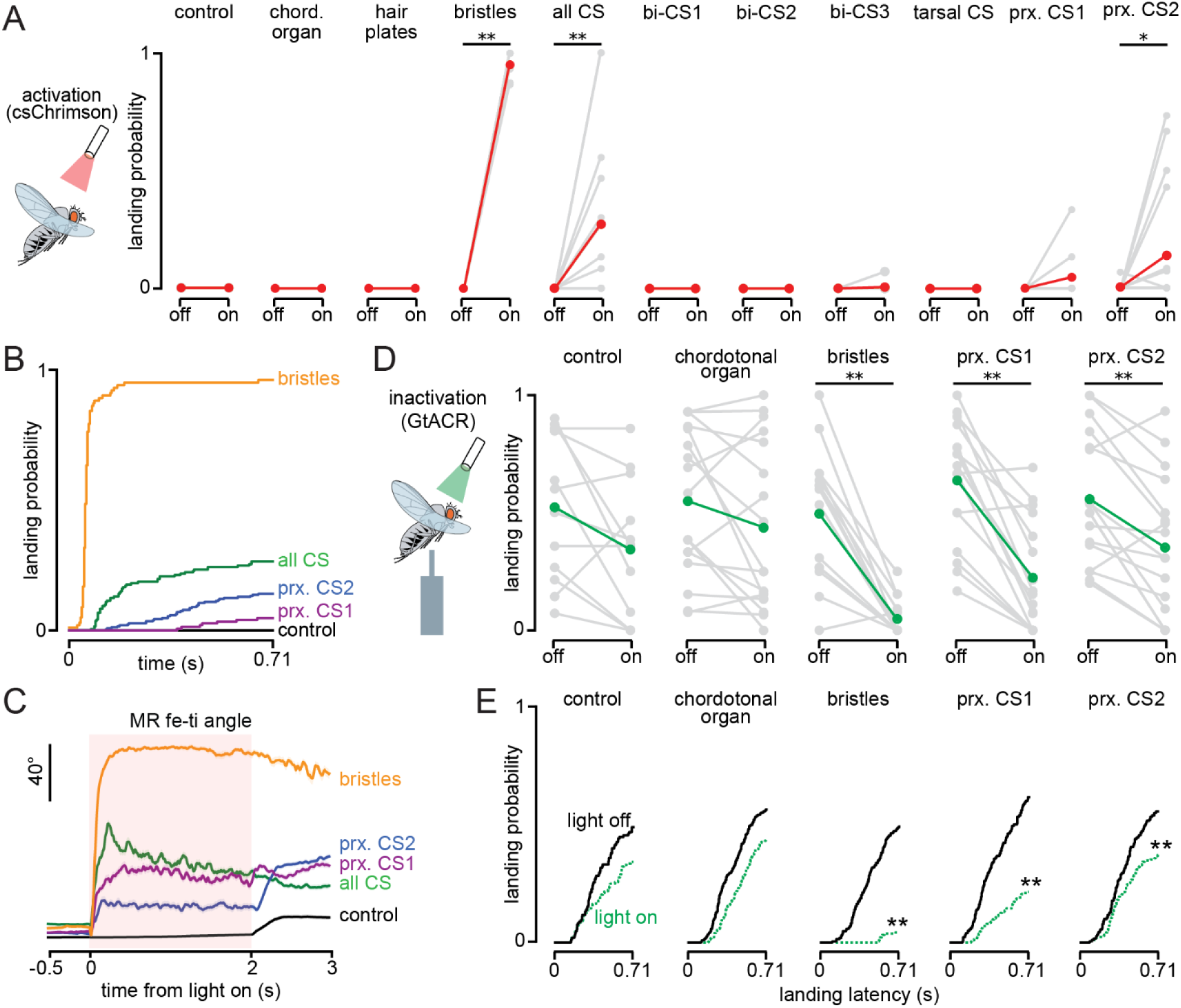
Optogenetic experiments confirm the role of tactile bristles and campaniform sensilla neurons in landing. **(A)** We used CsChrimson to optogenetically activate different subsets of mechanosensory neurons in tethered, flying flies. Fly-wise flight cessation probabilities are shown for light-ON (activated) and light-OFF (no activation) conditions in gray, along with overall population mean in red. From left to right, N=15, 12, 5, 7, 14, 5, 5, 13, 4, 10, and 18 flies. Statistical comparisons were performed using paired sign-flip permutation tests on fly-wise flight-cessation probability, \**p*<0.05, \*\**p*<0.01. Red light (660 nm) intensities: 0.0042 mWmm⁻² (control and all CS), 0.0423 mWmm⁻² (bristles), or 0.12 mWmm⁻² (all other groups). **(B)** Cumulative probability of flight cessation following optogenetic activation with CsChrimson. Statistical comparisons were performed using fly-wise restricted mean survival time (RMST) analysis with unpaired permutation tests across groups. Sample size is same as in (A). Black: control, green: all CS, brown: prx. CS1, blue: prx. CS2, and red: bristles. Exact *p* values and all pairwise comparisons are provided in Supplementary Table S2. **(C)** MR fe-ti joint angle aligned to light onset during optogenetic activation with csChrimson. Activation of bristles resulted in extension of the fe-ti joint angle. Activation of the different CS lines resulted in more moderate fe-ti extension. Solid lines indicate the mean and shaded areas indicate SEM. From top to bottom, n=60, 145, 209, 167, and 205 trials. **(D-E)** Optogenetic inactivation of mechanosensory neurons using GtACR. **(D)** Fly-wise landing probability under light on (inactivated) and light off (control) conditions is shown in gray, with the population mean shown in green. Green light (530 nm) was delivered at 0.121 mW mm⁻². Statistical comparisons were performed using paired sign-flip permutation tests on fly-wise landing probability, \*\**p*<0.01. From left to right, N=12, 17, 15, 17, and 17 flies. **(E)** Cumulative probability of landing following initial platform contact with light on (inactivate, green) or light off (control, black). Landing latency was analyzed up to the predefined landing threshold (τ = 0.71 s). Statistical comparisons were performed using fly-wise restricted mean survival time (RMST) analysis with paired sign-flip permutation tests. Sample sizes are the same as in (D), \*\**p*<0.01.

We found that activation of the different mechanosensory neurons also often led to other behavioral effects beyond flight cessation. For example, activation of proximal CS elicited grooming-like leg movements during flight, accompanied by moderate changes in femur-tibia joint angle (**Figure 5C**). Activation of tarsal bristles resulted in rapid and sustained leg extension (**Figure 5C**). As a result, we next tested whether these same sensory neurons are required for contact-mediated landing. We genetically expressed a light-gated chloride channel, GtACR1^21^, in mechanosensory neurons and then measured how landing probability in response to a mechanical contact changed when neurons were silenced. Inhibition of tactile bristle neurons and proximal CS significantly reduced landing likelihood and slowed landing latency, whereas inhibition of chordotonal organ neurons produced no detectable effect (**Figure 5D-E, Movie S3**). Thus, tactile bristles from the tarsus and proximally-located CS on the femur and trochanter are likely to underly contact-mediated landing, confirming the results of our connectivity analyses.

### An ascending neuron coordinates the transition from flight to landing

Landing is a complex behavior comprised of a sequence of sub-behaviors (**Figure 1A**)^5,8^. The timing of these behaviors and how the animal transitions from one to the next is key to a controlled and successful landing. Previous work found that looming visual input triggers an early landing behavior in flying flies, leg extension, via activation of various descending neurons including DNp10, DNp07, DNp47, and DNg79^22,23^ (**Figure 6A**). Our experiments examine the next step of landing, once the leg has made contact with the substrate and the animal maneuvers its legs so as to adhere and transition to a stable, standing posture and cease flying. At this stage of landing, the nervous system must rapidly transition from a visually-guided landing program, which promotes leg extension and body deceleration, to a contact-mediated motor program that stabilizes the body, establishes stance, and terminates flight. If the nervous system does not appropriately coordinate these behaviors, the fly may move its legs in the wrong direction and not successfully land. We therefore next sought to understand how neural circuits in the CNS facilitate this transition. We hypothesized the existence of a coordinating neuron or group of neurons that receive input from mechanosensory neurons detecting substrate contact and inhibit vision-mediated landing circuits, thereby facilitating the switch to grasping and flight termination (**Figure 6A**).

**Figure 6.**
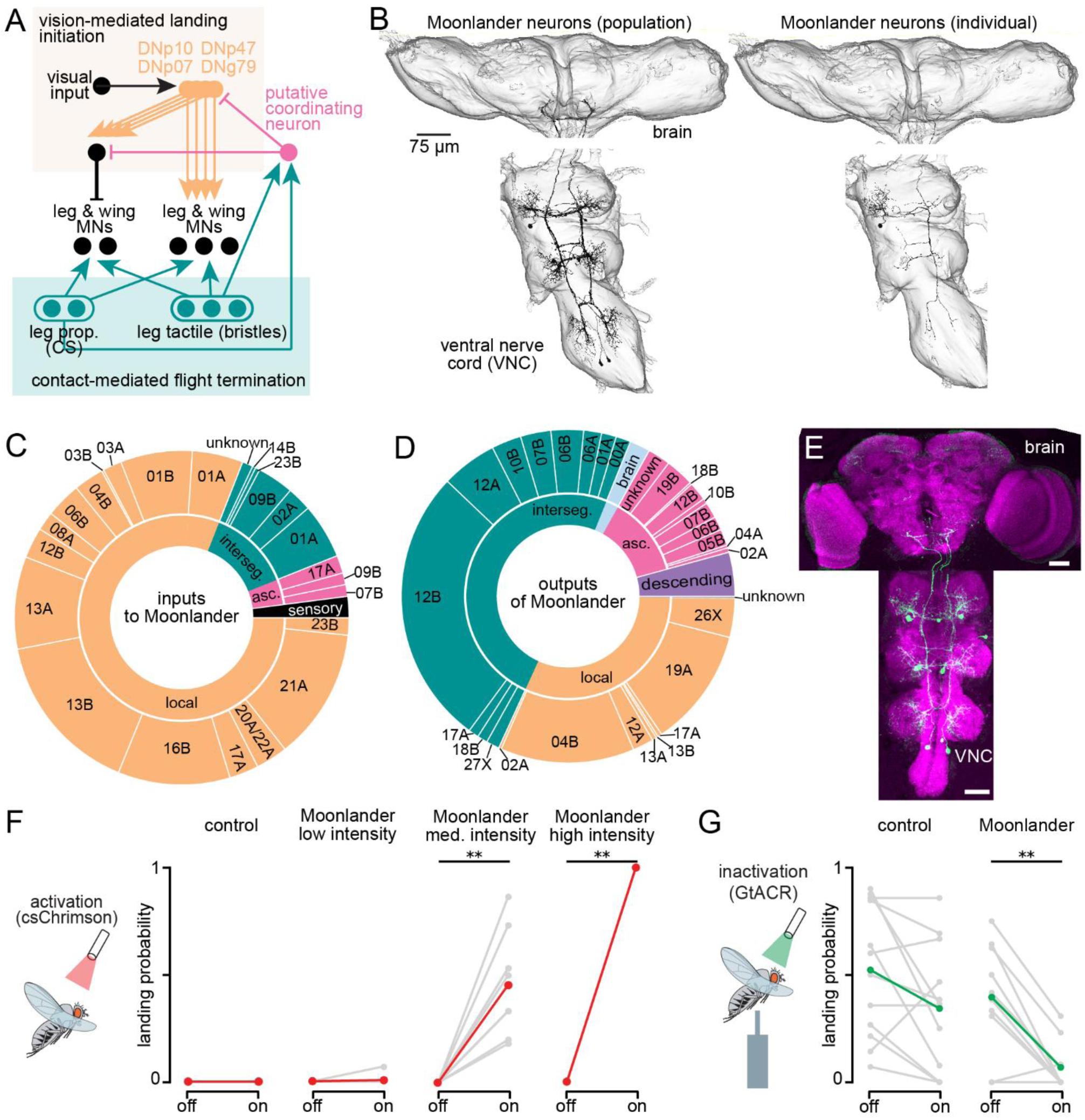
Moonlander ascending neurons mediate the switch from visually-induced landing initiation to contact-induced flight cessation. **(A)** We hypothesize the existence of a putative coordinating neuron (pink) that modulates the shift from vision-mediated landing initiation to touch-mediated flight cessation. **(B)** Connectomic analyses from BANC point to Moonlander neurons (AN06B002) as well-poised to play a coordinating role during landing. Left: image of all six Moonlander neurons in BANC. Right: a single Moonlander neuron with a cell body in the front left leg neuropil. **(C)** Moonlander neurons receive inputs from a variety of neurons, including leg mechanosensory neurons. **(D)** Moonlander neurons synapse onto a variety of brain and VNC neurons, including landing descending neurons identified by Ache et al (2019)^23^ and Liessem et al (2025)^22^. **(E)** Confocal images of genetically labeled Moonlander ascending neurons in the brain and central nerve cord (VNC). Magenta (nc82); green (GFP). Scale bar: 50 µm. **(F)** We optogenetically activated Moonlander ascending neurons in tethered flying flies via csChrimson. Fly-wise flight cessation probabilities are shown for light on (activated) and light off (no activation) conditions in gray, along with overall population mean in red. Red light (660 nm) was delivered at low (0.0042 mWmm⁻²), medium (0.0423 mWmm⁻²), or high (0.12 mWmm⁻²) intensities. From left to right, N=15, 14, 10, and 10 flies. Statistical comparisons were performed using paired sign-flip permutation tests on fly-wise flight-cessation probability, \*\**p*<0.01. **(G)** We optogenetically inactivated Moonlander ascending neurons in tethered flying flies via GtACR. Fly-wise landing probability under light on (inactivated) and light off (control) conditions is shown in gray, with the population mean in green. Green light (530 nm) was delivered at 0.121 mWmm⁻². From left to right, N=12 and 10 flies. Statistical comparisons were performed using paired sign-flip permutation tests on fly-wise landing probability, \*\**p*<0.01.

We again turned to connectomics to find neurons that are positioned to act as a coordinating node during landing. This time we used another recently published connectome, the BANC^24^, because it includes both the brain and VNC and would enable us to identify all ascending and descending neurons involved in the circuit in addition to those only within the VNC. We examined all neurons that receive input from any leg mechanosensory neurons and provide output onto the landing descending neurons identified by Liessem et al., (2025)^22^. Of these, the most strongly connected neuron that emerged was a group of 6 inhibitory GABAergic ascending neurons, AN06B002, hereafter referred to as “Moonlander” (**Figure 6B**). This cell type was previously described by Bates et al., (2026), who also hypothesized that it is likely involved in landing^24^. There is one Moonlander neuron per leg neuropil and they are all weakly interconnected with one another. Their inputs include leg mechanosensory neurons, ascending neurons, and VNC interneurons from a variety of hemilineages (**Figure 6C**). Intriguingly, ∼85% of the presynaptic partners of Moonlander neurons also receive input from leg mechanosensory neurons, implying strong direct and indirect connectivity with leg mechanosensory neurons. Moonlander neurons synapse onto several downstream partners, including the landing descending neuron, DNp10 (**Figure 6D**).

We developed a split-GAL4 line that labels Moonlander neurons (**Figure 6E**). We found that optogenetic activation of Moonlander ascending neurons in flying flies was sufficient to stop flight, and the likelihood of flight cessation depended on the light intensity and therefore intensity of activation (**Figure 6F, Movie S4**). Optogenetically inactivating these neurons, on the other hand, significantly decreased flies’ likelihood of landing, such that they rarely stopped flying even after being contacted by the motorized platform (**Figure 6G, Movie S4**). Together, our connectomics and optogenetic results identify Moonlander ascending neurons as a likely important node in the neural circuitry underlying landing and flight cessation. We propose that Moonlander neurons contribute to the transition from visually guided to contact mediated flight termination, potentially through modulation of landing-related descending neurons. Future experiments will be required to determine the precise circuit mechanisms by which Moonlander neurons coordinate this behavioral switch.

## Discussion

### Role of adaptive leg maneuvers in enabling successful landings

Flying animals have been shown to approach potential landing substrates via a variety of strategies that depend on the species, behavioral context, and substrate orientation^4–6,10,25–27^. Landing is thus a complex and heterogenous behavior, consisting of a series of sub-behaviors that must be flexible and yet coordinated in their sequence to enable robust performance. Sensory feedback, whether visual or mechanosensory, increases the envelope of landing success by coordinating these behaviors.

Towards this end, we found that small variations in mechanical contact can affect how the fly moves its legs and whether or not it lands and stops flying (**Figures 2-3**). Contact of the legs reliably induces landing, and contact of the distal ti-ta joint of the leg is more likely to lead to landing compared to the proximal cx-tr joint. Flies make directed leg movements that change in direction depending on where the mechanosensory contact is made, and additional contacts made by the other legs facilitate a faster and more probable landing and flight cessation. Thus, there is likely a complex set of neural circuits that enable the fly to determine the nature and location of any mechanical stimuli, including via exploration with its legs to make additional contacts, all with the goal of making a rapid decision of whether and how to land. Our experiments provide an entry point for mapping and understanding these circuits, and comparing them to those underlying other coordinated leg movements, like walking or grooming.

### Limitations of our behavioral assay

It is important to note that our behavioral assay does not completely recapitulate natural flying and landing. The flies in our assay are rigidly tethered and do not need to support their own weight, and they also have a more limited range of flight postures than would be possible during free flight. However, assaying the role of mechanosensory feedback in free flight experiments would be extremely challenging. Even though our flies were tethered and held in a stereotyped position, we still found it difficult to deliver a consistent mechanical stimulus. Small changes in the fly’s posture led to notably different post-contact movements of the contacted limb that affected the resulting behavior (**Figure 3**). This lack of stimulus stereotypy would be even greater during free flight. Thus, our experiments here provide a necessary starting point. Nevertheless, free flight experiments will still be essential to fully understand how the circuits and behaviors that we uncovered contribute to controlled landings. For example, the threshold we established for our tethered flight experiments, 0.71 s, enabled us to be inclusive in our analyses, but natural, free-flight landing transitions are typically quite rapid – as fast as 50 ms in some species^9^. It remains to be seen whether the searching leg movements that we describe here are as useful during such rapid maneuvers. Flies often perform rapid leg movements during takeoff^28^; similar movements are likely possible during landing. While the exact nature of the searching maneuvers may differ during untethered landings, coordinated mechanosensory feedback from multiple legs is likely to still play an important role in reinforcing the landing transition.

Another important distinction is that the flies in our assays are not actively planning to land when contact is made by the platform. While previous work has found that insects, including flies, can make unplanned and sudden landings, more often landings are planned and include a period of leg extension as the animals approach the landing substrate^26,29^. This posture has been shown to improve landing success by passively damping out collision forces^1^. However, it also almost certainly impacts how contact with the landing surface mechanically moves the leg, and therefore which mechanosensory neurons are activated. While we could not exactly mimic this interaction with our assay, we did find that contact of the distal leg joints so as to bend the leg outwards, similar to what typically happens during planned landings, increases landing probability and decreases landing latency (**Figure 3**). Thus, the neural circuits underlying landing may have evolved such that the contact that is most like the “expected” mechanosensory stimulus is most likely to lead to landing, whereas contacts that vary from this expectation may lead to additional leg movements and searching, delaying the termination of flight. This result suggests that landing is an active behavior in which the animal is continuously assessing substrate contact via mechanosensory feedback.

### Tactile bristles and CS mediate contact-initiated flight termination

Scientists have recognized for decades that surface contact is critical to mediating the transition to or from flight. Referred to as ‘Kontaktflugreflex’ or the tarsal reflex, most flying insects will initiate flight when their legs lose contact with a surface and stop flying when contact is regained^30,31^. Behavior, amputation experiments, and neurophysiology recordings have implicated tarsal contact bristles and leg campaniform sensilla as underlying this reflex^30–35^. Our results utilizing both connectomics and optogenetic experiments confirm these hypotheses (**Figures 4-5**). We are not yet able to determine the degree to which information from these two mechanosensory types inhibit flight via parallel or overlapping circuits. Our optogenetic experiments suggest that activating either tactile bristles or CS is sufficient to terminate flight, but also that inhibition of either is enough to interfere with landing (**Figure 5**).

Animals, including flies, have been shown to approach potential landing substrates via a variety of strategies that depend on substrate orientation. For example, when landing on inverted surfaces, flies initially accelerate upwards, rapidly rotate their bodies while extending their legs, and then perform a final leg-assisted swing after making contact with the substrate^36^. Landing strategies can also depend on the species. Bees frequently contact landing substrates first with their antennae^4^. Some species accelerate and crash into the landing substrate^26^ while mosquitoes do not appear to prepare for landing at all but rather depend on their forelimbs and proboscis to absorb the impact force^25^. These variations suggest that the neural circuits underlying sensory feedback during landing are likely species-specific and even context or substrate-specific. Thus, it is possible that different leg mechanosensory neurons are important for different landing maneuvers, even within *Drosophila.* In addition, while we found strong evidence for the role of the tarsal reflex in stopping flight, we rarely observed the complimentary reflex, that loss of contact with the platform would stimulate flight. As a result, we cannot determine if the mechanosensory neurons that mediate the flight to standing transition at landing are the same as those mediating the standing to flight transition at takeoff, as has been suggested in some species^30,31^. Based on connectomic analyses, Liessem et al., (2025) found that distinct neural circuits mediate landing and takeoff in *Drosophila*^22^. These behaviors may face distinct selection pressures despite seeming interrelated.

### The role of the CNS in coordinating landing-associated behaviors

Previous work studying landing has focused on the role of visual feedback in coordinating behaviors like approach, pitch, and leg extension^29,36,37^. Such feedback is important for ensuring that the animal is in the correct posture and position as to make a controlled landing. However, visual feedback is not the only source of sensory feedback, and it is likely less useful as the animal makes contact with the substrate, grasps on, and transitions to a stable standing posture. In addition, the leg and wing coordination patterns necessary for this touch-down phase of landing are likely distinct from those necessary for the approach phase^5,10,36^. Thus, as is the case with many complex behaviors, accomplishing a successful landing means the CNS must appropriately transition between sensorimotor control streams underlying distinct behavioral modules. Our experiments suggest that the fly does so via a sort of “behavioral switchboard” mediated by the Moonlander ascending neurons. These neurons receive input from leg mechanosensory neurons and synapse onto the previously identified landing descending neurons DNp10 and DNp07^22,23^. Inactivation of Moonlander neurons prevents the transition to standing and flight cessation after mechanical contact (**Figure 6**). Interestingly, even though Liessem et al., (2025) identified several other descending neurons that also receive visual input and are also involved in looming-induced landing initiation, Moonlander neurons only synapse onto DNp10 and DNp07^22^. We did not find any neurons that received direct input from leg mechanosensory neurons and synapsed onto the other landing neurons. This result may imply that mechanosensory feedback is only important during the subset of landing maneuvers mediated by DNp10 and DNp07, or perhaps that DNp10 and DNp07 are important nodes during all landing maneuvers, and that the other neurons identified by Liessem et al., simply further finetune the movements made during the approach phase. Nevertheless, we find strong evidence for a set of neural circuits responsible for sequencing the different sub-behaviors required for controlled landings. These circuits will be a useful model to understand how the CNS rapidly, precisely, and flexibly coordinates movements across multiple limbs to effect behavioral transitions.

## Acknowledgements

We thank Marie Suver, Chris Dallmann, Anthony Azevedo, and members of the Agrawal and Matthews lab for feedback on the manuscript. We thank Brandon Pratt, Anne Sustar, Leila Elabbady, John Tuthill, Anna Pierzchlińska, and Ansgar Büschges for providing fly stocks. We thank Brandon Pratt, Leila Elabbady, Shirin Mohammadian, Anthony Azevedo, Ellen Lesser, and Gwendolyn Swannell for neuron annotations from the FANC connectome. We thank Alexander Bates and Helen Yang for providing neuron annotations from the BANC connectome. This work was supported by NIH grant R00NS117657 to S.A..

## Author contributions

S.A. and T.K. conceived the project. S.A. acquired funding. T.K. performed all experiments and analyzed behavioral data. T.K., B.B., and B.F. annotated video frames to train our behavioral tracker. H.S. created and screened split Gal4 lines used in the project. F.B.G. contributed and analyzed confocal images. S.A. proofread and annotated neurons in FANC and BANC and analyzed all connectome data. T.K. and S.A. wrote the paper with input from all other co-authors.

## Data Availability

BANC connectome data presented in the paper was analyzed from the CAVE materialization v.888 (17 April 2026). FANC connectome data presented in the paper was analyzed from the CAVE materialization v604 timestamp 1684915801.222989. Annotated connectivity matrices are available as Python Pandas data frames (https://pandas.pydata.org/) at the GitHub repository: https://github.com/agrawallab3-hue/Kuo_landing_2026.

## Code Availability

Scripts to recreate the analyses and figures in the paper are available at Github: https://github.com/agrawallab3-hue/Kuo_landing_2026. All analysis was performed in Python 3.9 using custom code, making extensive use of CAVEclient (https://github.com/seung-lab/CAVEclient) and CloudVolume to interact with data infrastructure, and libraries Matplotlib, Numpy, Pandas, Scikit-learn, Scipy, stats-models, and VTK for general computation, machine learning, and data visualization.

## Declaration of generative AI and AI-assisted technologies

During the preparation of this work, the authors used OpenAI Codex to assist with code development, implementation of data analysis pipelines, debugging, and exploration of statistical analysis strategies, and ChatGPT (OpenAI) to assist with improving the clarity, organization, and language of the manuscript. The first draft was written by the authors.

## Methods

### Key resource table

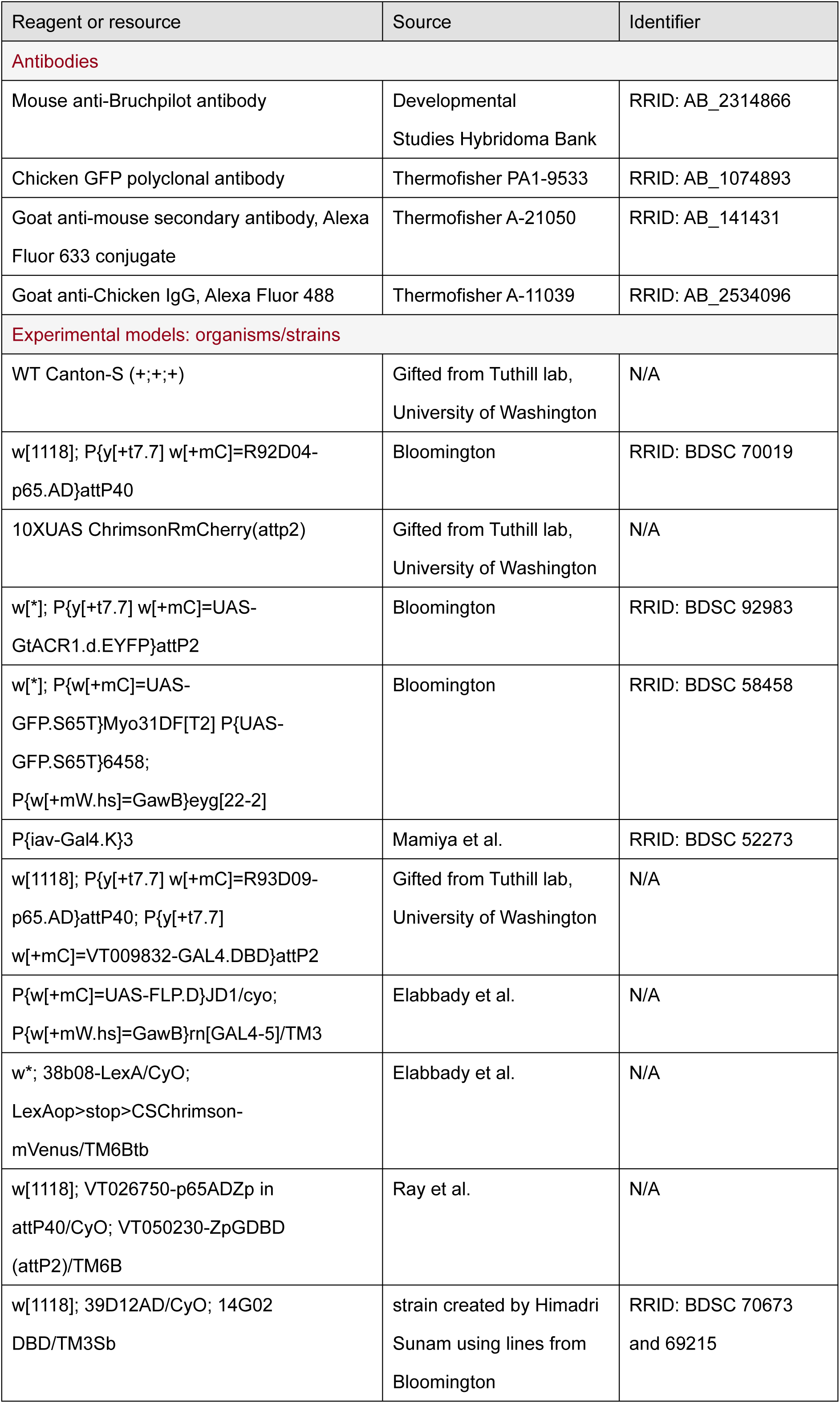
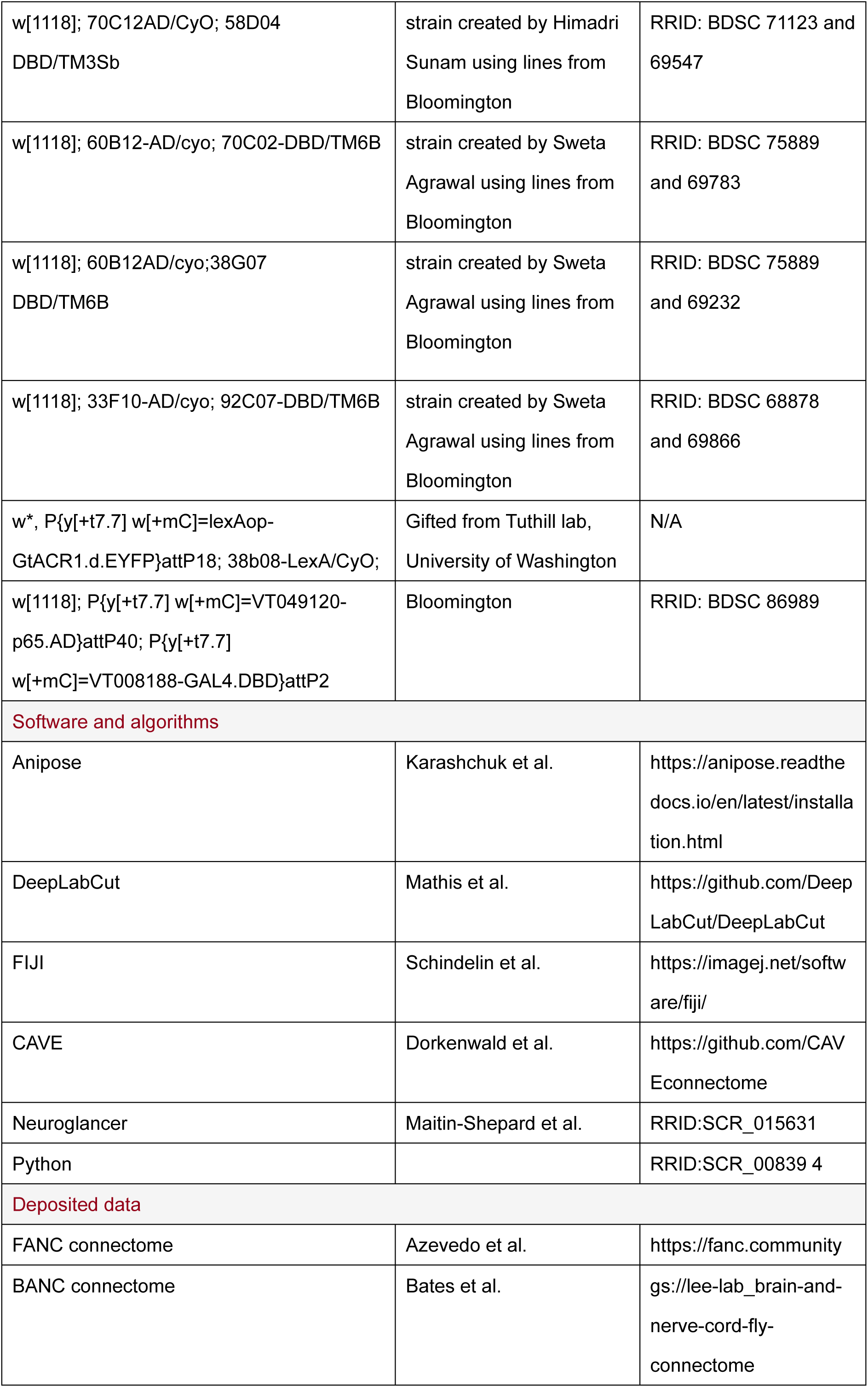

### Fly husbandry

*Drosophila melanogaster* were raised on standard cornmeal and molasses medium under a 14:10 hour light:dark cycle at 25°C. Female flies 3–5 days post-eclosion were used for all behavior experiments. For experiments involving optogenetic reagents, adult flies were placed on cornmeal agar with all-trans-retinal (35 mM in 95% EtOH; Santa Cruz Biotechnology) for approximately 48h prior to experiments. To prevent light exposure, vials were wrapped in aluminum foil during development. The genetic driver lines used for each experiment are listed in the resource table above.

### Fly preparation for *in vivo* optogenetics and landing experiments

Flies were transferred to a small 5 mL culture tube and cold anesthetized (<2 min). Flies were then transferred to a cold block for tethering (<10 min). The flies were glued to rigid tungsten tethers using ultraviolet-curing glue (Bondic) applied to the rostral dorsal thorax to ensure mobility of wings during flight.

### Behavior assay

Tethered flies were suspended in midair in a completely dark behavioral chamber. To elicit landing behavior, custom-built platforms made of aluminum tubing and tungsten wire were attached to a linear actuator (3574_0, Phidget Inc). A PID controller (custom-written code) was used to control the platform’s movement towards the fly. Due to considerations surrounding the accessibility of different leg joints during flight, we used two different platform probes. The tibia-tarsus (ti-ta) joint was more variable in its position during flight and also more prone to slipping away from the probe, and so we contacted it using a wider probe. In contrast, we used a narrower probe to contact the coxa-trochanter (cx-tr) joint so as to avoid contacting any other leg joints. Each trial lasted 7 s with a 10 s inter-trial interval. The platform reached its target position 3 s after trial start, remained in position for 1 s, and was then retracted.

A gentle air puff aimed at the fly was delivered at 5 s after the trial end and prior to the next trial start to re-initiate flight during the inter-trial interval. This air puff was delivered regardless of whether the fly had stopped flying. For conventional landing experiments, each fly experienced 20 trials of repeated mechanical perturbation. For optogenetic experiments, 30 trials were conducted per fly, consisting of 15 light on and 15 light off trials in randomized order. For activation experiments involving ChrimsonR, a red light with low (0.0042mW/mm^2^), medium (0.0423 mW/mm^2^) or high (0.12mW/mm^2^) intensity with wavelength of 660 nm was delivered as a 30 Hz pulse train with a 22-ms pulse width (66% duty cycle) for 2 s beginning 3 s after trial onset. For these trials, no mechanical perturbation was presented to the fly. For inactivation experiments with GtACR, a green light (0.121 mW/mm^2^, 530 nm) was delivered as a 30 Hz pulse train with a 22-ms pulse width (66% duty cycle) throughout the full 7 s trial, including during platform contact.

### Kinematics tracking

Tethered flies were surrounded by six cameras (aca800-510um, Basler) recording throughout each trial. The fly was illuminated by two infrared (IR) light sources (LM45, Smart Vision, 850 nm). During video acquisition, we strobed the IR lights for 80 µs synchronized to the start of each frame capture to reduce the motion blur of the fly’s limbs. Acquisition parameters differed slightly depending on experiment type, due to continual improvements made to the recording and tracking pipeline. Wild-type mechanical contact experiments (**Figure 2B**) were recorded for 7 s at 200-250 Hz. The no-contact group was recorded at 300 Hz (n=10). All optogenetic activation and inactivation experiments were recorded at 250 Hz. Ablation experiments (**Figure 2K**) were performed earlier in the study and were recorded at 300 Hz without IR strobing.

A subset of recorded frames from each camera were extracted and labeled using the image annotator Anivia (https://lambdaloop.com/anivia/). We used these annotations to train a deep neural network (DeepLabCut^38^) to automatically track all leg joints in each camera view. Predictions with less than 0.9 confidence scores were filtered out. 2D tracking data from all six camera views were then combined to reconstruct leg joint positions and angles in 3D using Anipose^39^.

### Quality Control of Kinematic Tracking

Tracking quality was evaluated at the frame level for each keypoint used in the analysis. A frame was marked invalid if the reconstructed 3D coordinate was missing, the reprojection error was missing or exceeded the preset threshold of 50, the camera count was missing or below the minimum required number of two cameras, or the tracking score/likelihood was missing or below threshold of 0.8. Invalid samples were set to missing values, and short invalid gaps were linearly interpolated only when the gap length was shorter than 20 ms. For each analysis window, the invalid frame fraction was calculated as the number of invalid frames divided by the total number of frames in that window. Trials were excluded if the invalid frame fraction exceeded the preset tolerance of 30% or if the longest invalid gap exceeded 20 ms.

Joint angles were calculated from 3 reconstructed 3D landmarks, with the middle landmark defining the vertex of the angle. For angle analyses, tracking quality control (QC) was applied to each of the three landmarks defining the joint angle before angle calculation. Each landmark trace was filtered and short invalid gaps were interpolated using the same QC criteria described above. A trial was retained only if all three landmark traces passed QC within the analysis window. Joint angles were then calculated from the QC-cleaned 3D coordinates, with the middle landmark defining the angle vertex, and the resulting angle traces were smoothed with an exponential moving average before averaging and visualization.

### Behavioral analysis

Trials were excluded from analysis if flies were not actively flying when the platform made contact or if the platform failed to contact the target joints properly. Improper contacts included absence of platform contact, slipping legs during contact, or contact with incorrect body parts. Flies were excluded from further analysis if fewer than half of the total trials were valid. For non-optogenetic experiments, which consisted of 20 trials per fly, flies were excluded if more than 10 trials were invalid. For optogenetic experiments, flies were excluded if more than 7 trials were invalid in either the light on or light off condition, which consists of 15 trials each. For all subsequent analyses except optogenetic activation and inactivation experiments, the first 3 trials were excluded because flies exhibited noticeable changes in flight posture during these initial trials.

Landing probability was calculated only after trial and fly exclusion. Specifically, flies first had to satisfy the inclusion criteria described above. A trial was classified as “landing” when the targeted body parts successfully made the first contact with the platform followed by flight cessation within 0.71s after platform contact. Trials in which flight cessation occurred after this threshold or did not occur within the observation window were classified as “flying” trials.

The 0.71-s landing threshold was determined from the raw landing latency distribution of the right middle tibia–tarsus contact (MR ti-ta) dataset after excluding the first three trials. Because the latency distribution exhibited moderate skew, the threshold was estimated as the median plus two median absolute deviations (MAD×1.483), where the scaling factor (1.4286) approximates the standard deviation for normally distributed data, providing an analogue to the conventional mean + 2 SD criterion. This threshold was used to determine whether a flight cessation event could reasonably be attributed to mechanical perturbation. Flies were not considered to have landed if they stopped flying without folding their wings or if flight cessation occurred after the predetermined latency threshold.

### Subsequent Leg Contact Timing and Survival Analysis

Subsequent leg contact (SLC) latency was defined as the earliest time point at which any of the non-contacted legs made stable contact with the platform following the initial contact of the targeted leg. To ensure that SLC events were behaviorally relevant to the landing response, only events occurring within the analysis window were considered valid. For successful landing trials, valid SLC events were required to occur between the moment of contact (MOC) and moment of landing (MOL), which is determined by flight cessation. For unsuccessful trials, valid SLC events were required to occur between MOC and the predetermined landing latency threshold (0.71 s). For the per-leg SLC-latency analyses, trials in which a given leg did not establish a valid SLC within the corresponding analysis window were treated as censored for that leg (**Figure 2G**). Separately, for first-SLC analyses, the earliest valid SLC across all non-contacted legs was identified for each trial and the identity of the contacting leg was recorded. Fly-wise first-SLC probability was then calculated for each contralateral non-contacted leg as the proportion of valid trials in which that leg was the first to establish a subsequent contact. (**Figure 2H**)

### Statistical Analyses

Unless otherwise specified, the individual fly was treated as the independent biological replicate and statistical unit. Repeated trials from the same fly were therefore summarized at the fly level before statistical comparison. Individual trials were treated as independent observations only for analyses explicitly described as trial-level, including the SLC-count analysis in **Figure 2J** and the analyses in **Figure 3** described below. All statistical tests were two-sided.

#### Permutation tests of fly-wise probabilities

Landing and flight-cessation probabilities were calculated separately for each fly after applying all trial and fly exclusion criteria. First subsequent leg contact (SLC) probabilities were similarly calculated separately for each fly as described above. For comparisons between independent groups (e.g., different mechanical stimulation conditions), fly-wise probabilities were compared using two-sided unpaired permutation tests on the difference in group means. For paired comparisons (e.g., light on versus light off conditions in optogenetic experiments), fly-wise probabilities were compared using paired sign-flip permutation tests on the within-fly probability differences. These analyses were used for landing probability, first subsequent leg contact (SLC) probability, and optogenetic landing and flight-cessation probability analyses.

#### Permutation tests of fly-wise restricted mean survival time

Landing latency and subsequent leg contact (SLC) latency were analyzed as right-censored time-to-event data. Successful landing or SLC events were treated as events, whereas trials without an event within the observation window were treated as censored. For each fly and experimental condition, latency distributions were summarized using the restricted mean survival time (RMST) up to a fixed time horizon (τ = 0.71 s).

Comparisons between independent groups were performed using two-sided unpaired permutation tests on fly-wise RMST values, whereas paired comparisons were performed using paired sign-flip permutation tests on within-fly RMST differences.

Trial-level analyses of behavior-defined groups Analyses in **Figure 3** treated individual trials as independent observations. For contact-geometry analyses, including transient touch (TT) versus sustained touch (ST), inward touch (IT) versus outward touch (OT), trials were compared using two-sided trial-level label-shuffle permutation tests on binary landing outcomes. Each trial was assigned a value of 1 for successful landing and 0 for unsuccessful landing. The observed statistic was the difference in mean landing probability between behavioral groups. Trial labels were randomly shuffled while preserving the original group sizes, and statistical significance was determined from the permutation distribution. Landing-latency distributions between behavioral trial groups were analyzed using Kaplan–Meier survival analysis with right censoring. Curves were plotted as inverted survival functions (1 − S(t)), and differences between behavioral groups were assessed using two-sided log-rank tests.

For path efficiency analyses, successful and no landing trials were compared using two-sided unpaired permutation test on trial-level path efficiency values. The association between trial-level path efficiency and landing latency were assessed using spearman’s rank correlation. Spearman’s correlation coefficient (ρ) was reported as the measure of association.

#### Trial-level SLC-count analysis

For the analysis of subsequent leg contact (SLC) number in **Figure 2J**, individual trials were treated as independent observations. The number of valid SLC events occurring within the predefined behavioral analysis window was calculated for each trial. Successful and unsuccessful landing trials were compared using a two-sided unpaired permutation test on trial-level SLC counts, with the difference in mean SLC count between groups as the test statistic. Trial labels were randomly shuffled between landing-outcome groups while preserving the original group sizes.

#### Fly-wise displacement-vector analysis

For analysis of projected ti-ta trajectories, endpoint displacement in each trial was represented as a two-dimensional vector from the ti-ta position at the moment of contact to the selected trajectory endpoint. Displacement vectors were averaged across trials within each fly and contact group, with the fly treated as the statistical unit. For the primary analysis, the direction of each fly-mean displacement vector was expressed as an angle calculated from its two-dimensional vector components. Directional differences between contact groups were assessed using two-sided fly-wise directional displacement unpaired permutation tests with the angular distance between the circular mean directions of the two groups as the test statistic. Statistical significance was determined by permuting fly-level group labels while preserving the original group sizes. (**Figure 2I**) As a secondary analysis, differences in the full two-dimensional displacement vectors between contact groups were assessed using two-sided fly-wise two dimensional permutation tests, with Euclidean distance between group mean displacement vectors as the test statistic. Statistical significance was determined using the same fly-level label shuffle procedure. Additional secondary analyses separately compared the x component, y component, and magnitude of the fly-mean displacement vectors using two-sided unpaired permutation test at the fly level. (**Figure 2F**)

#### Permutation test

Unless otherwise specified, permutation tests were performed by randomly shuffling group labels while preserving the original sample sizes. For analysis in which the fly was the statistical unit, group labels were permuted at the fly level. For trial-level permutation analyses in **Figures 2J and 3**, labels were permuted at the trial level. For paired comparisons, permutation testing was performed by randomly flipping the sign of within-fly differences. *P* values were calculated from 20,000 permutations as the proportion of randomized test statistics whose absolute value was greater than or equal to the observed statistic with + 1 correction.

#### Multiple-comparison correction

For analyses involving multiple pairwise comparisons addressing the same predefined statistical question, those analyses were treated as a single comparison family, and adjusted P values were calculated as P_adj_ = min(P_raw_×m, 1), where m is the number of comparisons within that family. The composition and number of comparisons in each statistical family, together with raw and adjusted P values, are reported in the corresponding Supplementary Tables. S[1][2]

### Immunohistochemistry and confocal imaging

We drove the expression of mCD8-GFP in neurons labeled by each GAL4 driver line. We then examined expression in the VNC after fixing in a 4% paraformaldehyde (PFA) PBS solution for 20 minutes and incubating with first primary antibodies (anti-GFP chicken polyclonal antibody; anti-brp mouse for nc82 neuropil staining) overnight and then secondary antibodies (anti-chicken-Alexa 488; anti-mouse-Alexa 633) overnight. Using a Zeiss LSM900 confocal microscope, we imaged the expression of GFP in the VNC. We used FIJI^40^ to post-process all confocal stacks.

### Reconstruction and identification of leg mechanosensory axons in FANC

We obtained hair plate neuron, femoral chordotonal organ neuron, tactile bristle axon, wing motor neuron, and wing pre-motor hemilineage annotations from already published data^13,17,20,41^. We reconstructed campaniform sensilla axons in the electron microscopy dataset of the female ventral nerve cord, FANC, through manual proofreading of the automatically segmented cell fragments in Google’s collaborative Neuroglancer interface^42^. Neuron annotations were managed by CAVE, the Connectome Annotation Versioning Engine^43^. We used custom Python scripts to interact with CAVE via CAVEclient. Via these scripts, we queried all monosynaptic and disynaptic connections between leg mechanosensory neurons and wing motor neurons that had a connection strength of 10 synapses or more. All interneurons were typed according to hemilineage by Lesser et al., (2024)^13^ and then verified by us via visual inspection.

### Reconstruction and identification of landing-related central neurons in BANC

We obtained annotations for all leg mechanosensory neurons, landing descending neurons, and interneuron hemilineages from already published data^24^, with an additional verification step performed by us via visual inspection. We queried all neurons that received at least 4 synapses from at least one leg mechanosensory neuron and made at least 4 synapses onto at least one descending landing-initiation neuron as identified by Liessem et al (2025)^22^. Of these, AN06B002 neurons were among the most strongly connected to both leg mechanosensory neurons and landing-initiation descending neurons. We then queried and classified all neurons that made or received at least 20 synapses cumulative onto any of the AN06B002 neurons. Neuron hemilineages were either taken from already published data^24^ or determined via visual inspection.

