## supplemental legends, tables 1 and 2 for "Sticking the landing: leg mechanosensory feedback coordinates the transition from flight to landing"

**Movie S1: Mechanical stimulation of the coxa-trochanter or femur-tibia joints of the middle leg.** Videos of tethered wild-type (*canton-S*) flies as platform is raised to contact the right middle-leg (MR) coxa–trochanter (cx-tr) or tibia–tarsus (ti-ta) joint. The displayed timer indicates time relative to the moment of platform contact, with 0 s corresponding to initial platform contact with the targeted joint.

**Movie S2: Optogenetic activation of mechanosensory neurons in tethered flying flies.** Videos of tethered flying flies during CsChrimson-mediated activation of control (92D04-AD), proximal campaniform sensilla 1 (prx. CS1), proximal campaniform sensilla 2 (prx. CS2), all campaniform sensilla (all CS), and tarsal bristle groups. The red dot indicates light onset. The displayed timer indicates time relative to red light onset.

**Movie S3: Mechanical stimulation of the femur-tibia joint of the middle leg during optogenetic inactivation of mechanosensory neurons.** Videos of tethered flying flies during GtACR-mediated inactivation of control (empty-GAL4), proximal campaniform sensilla 1 (prx. CS1), proximal campaniform sensilla 2 (prx. CS2), femoral chordotonal organ (FeCO), and tarsal bristle groups. The green dot indicates light onset. The displayed timer indicates time relative to the moment of platform contact, with 0 s corresponding to initial platform contact.

**Movie S4: Optogenetic activation and inactivation of Moonlander ascending neurons.** Videos of tethered flying flies during optogenetic activation or inactivation of Moonlander ascending neurons. For activation experiments, the videos illustrate behavioral responses evoked by CsChrimson-mediated activation; the red dot indicates light onset, and the displayed timer indicates time relative to light onset. For inactivation experiments, the videos illustrate behavioral responses to mechanical stimulation during GtACR-mediated inactivation; the green dot indicates light onset, and the displayed timer indicates time relative to the moment of platform contact.

**Table S1. Comprehensive statistical comparisons and multiple-comparison adjustments related to Figure 2:** The statistical analysis is specified at the beginning of each block. Unless otherwise specified, the individual fly is the independent biological replicate and statistical unit. Across-group comparisons use two-sided unpaired permutation tests; within-fly comparisons use paired sign-flip permutation tests. All permutation tests use 20,000 permutations with a +1 correction. For each predefined comparison family, Bonferroni-adjusted P values were calculated as $P_{adj}=\min(P_{raw}\times m, 1)$, where m is shown in the block note. Significance notation: ns: $P_{adj}\geq 0.05$; *: $P_{adj}<0.05$; **: $P_{adj}<0.01$; ***: $P_{adj}<0.001$.

**Figure 2A: Fly-wise landing probability.** Two-sided unpaired permutation test on fly-wise landing probability. Main joint-comparison family Bonferroni family m = 9.

| Comparison | N (flies) | Δ probability | $P_{raw}$ | $P_{adj}$ | Sig. |
|---|---|---|---|---|---|
| FR-ti-ta vs. MR-ti-ta | 15/14 | 0.09467 | 0.267 | 1 | ns |
| FR-ti-ta vs. HR-ti-ta | 15/14 | 0.16503 | 0.029 | 0.261 | ns |
| MR-ti-ta vs. HR-ti-ta | 14/14 | 0.07035 | 0.400 | 1 | ns |
| FR-cx-tr vs. MR-cx-tr | 12/17 | -0.03782 | 0.169 | 1 | ns |
| FR-cx-tr vs. HR-cx-tr | 12/17 | -0.00319 | 0.932 | 1 | ns |
| MR-cx-tr vs. HR-cx-tr | 17/17 | 0.03462 | 0.255 | 1 | ns |
| FR-ti-ta vs. FR-cx-tr | 15/12 | -0.68454 | $5\times10^{-5}$ | $4.5\times10^{-4}$ | *** |
| MR-ti-ta vs. MR-cx-tr | 14/17 | -0.81703 | $5\times10^{-5}$ | $4.5\times10^{-4}$ | *** |
| HR-ti-ta vs. HR-cx-tr | 14/17 | -0.85277 | $5\times10^{-5}$ | $4.5\times10^{-4}$ | *** |

**Figure 2A: Control comparisons.** Two-sided unpaired permutation test on fly-wise landing probability. Control-comparison family Bonferroni family m = 3.

| Comparison | N (flies) | Δ probability | $P_{raw}$ | $P_{adj}$ | Sig. |
|---|---|---|---|---|---|
| No contact vs. Abdomen | 10/11 | -0.02222 | 0.243 | 0.729 | ns |
| No contact vs. Abd w/o HL | 10/9 | -0.00714 | 1 | 1 | ns |
| Abdomen vs. Abd w/o HL | 11/9 | -0.02936 | 0.069 | 0.207 | ns |

**Figure 2B: Fly-wise landing latency (RMST).** Two-sided unpaired permutation test on fly-wise RMST ($\tau$ = 0.71 s). Bonferroni family m = 9.

| Comparison | N (flies) | Δ RMST | $P_{raw}$ | $P_{adj}$ | Sig. |
|---|---|---|---|---|---|
| FR-ti-ta vs. MR-ti-ta | 15/14 | -0.06227 | 0.202 | 1 | ns |
| FR-ti-ta vs. HR-ti-ta | 15/14 | -0.10733 | 0.023 | 0.207 | ns |
| MR-ti-ta vs. HR-ti-ta | 14/14 | -0.04505 | 0.408 | 1 | ns |
| FR-cx-tr vs. MR-cx-tr | 12/17 | 0.01015 | 0.198 | 1 | ns |
| FR-cx-tr vs. HR-cx-tr | 12/17 | 0.00721 | 0.408 | 1 | ns |
| MR-cx-tr vs. HR-cx-tr | 17/17 | -0.00293 | 0.653 | 1 | ns |
| FR-ti-ta vs. FR-cx-tr | 15/12 | 0.25213 | $5\times10^{-5}$ | $4.5\times10^{-4}$ | *** |
| MR-ti-ta vs. MR-cx-tr | 14/17 | 0.32456 | $5\times10^{-5}$ | $4.5\times10^{-4}$ | *** |
| HR-ti-ta vs. HR-cx-tr | 14/17 | 0.36668 | $5\times10^{-5}$ | $4.5\times10^{-4}$ | *** |

**Figure 2F: Fly-wise two-dimensional endpoint displacement.** Two-sided fly-wise 2D displacement-vector permutation test; P values are based on Euclidean distance between group-mean vectors.

Bonferroni family m = 9.

| Comparison | N (flies) | Vector difference | $P_{raw}$ | $P_{adj}$ | Sig. |
|---|---|---|---|---|---|
| FR-ti-ta-FL vs. MR-ti-ta-FL | 15/14 | Δx = 0.11858; Δy = -0.20668 | $6.5\times10^{-4}$ | $5.85\times10^{-3}$ | ** |
| FR-ti-ta-FL vs. HR-ti-ta-FL | 15/14 | Δx = -0.2541; Δy = -0.02066 | $5\times10^{-5}$ | $4.5\times10^{-4}$ | *** |
| MR-ti-ta-FL vs. HR-ti-ta-FL | 14/14 | Δx = -0.37269; Δy = 0.18603 | $1\times10^{-4}$ | $9\times10^{-4}$ | *** |
| FR-ti-ta-HL vs. MR-ti-ta-HL | 15/14 | Δx = 0.00858; Δy = -0.14785 | 0.066 | 0.594 | ns |
| FR-ti-ta-HL vs. HR-ti-ta-HL | 15/14 | Δx = 0.54619; Δy = 0.08420 | $5\times10^{-5}$ | $4.5\times10^{-4}$ | *** |
| MR-ti-ta-HL vs. HR-ti-ta-HL | 14/14 | Δx = 0.53761; Δy = 0.23205 | $5\times10^{-5}$ | $4.5\times10^{-4}$ | *** |
| FR-ti-ta-ML vs. MR-ti-ta-ML | 15/14 | Δx = 0.10342; Δy = -0.19344 | $5\times10^{-5}$ | $4.5\times10^{-4}$ | *** |
| FR-ti-ta-ML vs. HR-ti-ta-ML | 15/14 | Δx = 0.25962; Δy = -0.34572 | $5\times10^{-5}$ | $4.5\times10^{-4}$ | *** |
| MR-ti-ta-ML vs. HR-ti-ta-ML | 14/14 | Δx = 0.1562; Δy = -0.15229 | $1\times10^{-4}$ | $9\times10^{-4}$ | *** |

**Figure 2G: SLC latency (RMST), across stimulation groups.** Two-sided unpaired permutation test on fly-wise SLC RMST. Bonferroni family m = 12.

| Comparison | N (flies) | Δ RMST | $P_{raw}$ | $P_{adj}$ | Sig. |
|---|---|---|---|---|---|
| FR-ti-ta-FL vs. MR-ti-ta-FL | 15/14 | 0.12304 | 0.029 | 0.348 | ns |
| FR-ti-ta-HL vs. MR-ti-ta-HL | 15/14 | -0.16783 | 0.003 | 0.036 | * |
| FR-ti-ta-ML vs. MR-ti-ta-ML | 15/14 | -0.08849 | 0.089 | 1 | ns |
| FR-ti-ta-HR vs. MR-ti-ta-HR | 15/14 | -0.15488 | 0.002 | 0.024 | * |
| FR-ti-ta-FL vs. HR-ti-ta-FL | 15/14 | 0.33218 | $5\times10^{-5}$ | $6\times10^{-4}$ | *** |
| FR-ti-ta-HL vs. HR-ti-ta-HL | 15/14 | -0.19489 | $3\times10^{-4}$ | $3.6\times10^{-3}$ | ** |
| FR-ti-ta-ML vs. HR-ti-ta-ML | 15/14 | 0.26836 | $5\times10^{-5}$ | $6\times10^{-4}$ | *** |
| FR-ti-ta-MR vs. HR-ti-ta-MR | 15/14 | 0.25615 | $5\times10^{-5}$ | $6\times10^{-4}$ | *** |
| MR-ti-ta-FL vs. HR-ti-ta-FL | 14/14 | 0.20913 | $5\times10^{-5}$ | $6\times10^{-4}$ | *** |
| MR-ti-ta-HL vs. HR-ti-ta-HL | 14/14 | -0.02706 | 0.632 | 1 | ns |
| MR-ti-ta-ML vs. HR-ti-ta-ML | 14/14 | 0.35686 | $5\times10^{-5}$ | $6\times10^{-4}$ | *** |
| MR-ti-ta-FR vs. HR-ti-ta-FR | 14/14 | 0.17085 | $5\times10^{-5}$ | $6\times10^{-4}$ | *** |

**Figure 2G: SLC latency (RMST), within FR-ti-ta group.** Paired sign-flip permutation test on within-fly SLC RMST differences. Bonferroni family m = 10.

| Comparison | N (flies) | Δ RMST | $P_{raw}$ | $P_{adj}$ | Sig. |
|---|---|---|---|---|---|
| FL vs. ML | 15 | 0.06382 | 0.008 | 0.08 | ns |
| FL vs. HL | 15 | 0.19411 | $1\times10^{-4}$ | $1\times10^{-3}$ | *** |
| FL vs. MR | 15 | -0.01857 | 0.115 | 1 | ns |
| FL vs. HR | 15 | 0.1506 | $2\times10^{-4}$ | $2\times10^{-3}$ | ** |

| Comparison | N (flies) | Δ RMST | $P_{raw}$ | $P_{adj}$ | Sig. |
|---|---|---|---|---|---|
| ML vs. HL | 15 | 0.13029 | $5\times10^{-5}$ | $5\times10^{-4}$ | *** |
| ML vs. MR | 15 | -0.08239 | $1\times10^{-4}$ | $1\times10^{-3}$ | *** |
| ML vs. HR | 15 | 0.08678 | $2\times10^{-4}$ | $2\times10^{-3}$ | ** |
| HL vs. MR | 15 | -0.21268 | $5\times10^{-5}$ | $5\times10^{-4}$ | *** |
| HL vs. HR | 15 | -0.04351 | 0.033 | 0.33 | ns |
| MR vs. HR | 15 | 0.16917 | $2\times10^{-4}$ | $2\times10^{-3}$ | ** |

**Figure 2G: SLC latency (RMST), within MR-ti-ta group.** Paired sign-flip permutation test on within-fly SLC RMST differences. Bonferroni family m = 10.

| Comparison | N (flies) | Δ RMST | $P_{raw}$ | $P_{adj}$ | Sig. |
|---|---|---|---|---|---|
| FL vs. ML | 14 | -0.14772 | $3\times10^{-4}$ | 0.003 | ** |
| FL vs. HL | 14 | -0.09677 | $6.5\times10^{-4}$ | $6.5\times10^{-3}$ | ** |
| FL vs. FR | 14 | 0.03829 | 0.158 | 1 | ns |
| FL vs. HR | 14 | -0.12733 | 0.002 | 0.02 | * |
| ML vs. HL | 14 | 0.05095 | 0.001 | 0.01 | ** |
| ML vs. FR | 14 | 0.18601 | $5\times10^{-5}$ | $5\times10^{-4}$ | *** |
| ML vs. HR | 14 | 0.02039 | 0.283 | 1 | ns |
| HL vs. FR | 14 | 0.13506 | 0.001 | 0.01 | ** |
| HL vs. HR | 14 | -0.03056 | 0.173 | 1 | ns |
| FR vs. HR | 14 | -0.16562 | $9.5\times10^{-4}$ | $9.5\times10^{-3}$ | ** |

**Figure 2G: SLC latency (RMST), within HR-ti-ta group.** Paired sign-flip permutation test on within-fly SLC RMST differences. Bonferroni family m = 10.

| Comparison | N (flies) | Δ RMST | $P_{raw}$ | $P_{adj}$ | Sig. |
|---|---|---|---|---|---|
| FL vs. ML | 14 | 0 | 1 | 1 | ns |
| FL vs. HL | 14 | -0.33296 | $1.5\times10^{-4}$ | $1.5\times10^{-3}$ | ** |
| FL vs. FR | 14 | 0 | 1 | 1 | ns |
| FL vs. MR | 14 | -0.09461 | 0.002 | 0.02 | * |
| ML vs. HL | 14 | -0.33297 | $1.5\times10^{-4}$ | $1.5\times10^{-3}$ | ** |
| ML vs. FR | 14 | 0 | 1 | 1 | ns |
| ML vs. MR | 14 | -0.09461 | 0.002 | 0.02 | * |
| HL vs. FR | 14 | 0.33296 | $2\times10^{-4}$ | $2\times10^{-3}$ | ** |
| HL vs. MR | 14 | 0.23836 | $2\times10^{-4}$ | $2\times10^{-3}$ | ** |
| FR vs. MR | 14 | -0.09461 | 0.002 | 0.02 | * |

**Figure 2H: First-SLC probability, within stimulation groups.** Paired sign-flip permutation test on within-fly first-SLC probability differences. Bonferroni family m = 9.

| Comparison | N (flies) | Δ probability | $P_{raw}$ | $P_{adj}$ | Sig. |
|---|---|---|---|---|---|
| FR-ti-ta-FL vs. FR-ti-ta-ML | 15 | -0.38095 | $4.8\times10^{-3}$ | 0.043 | * |
| FR-ti-ta-FL vs. FR-ti-ta-HL | 15 | -0.60843 | $1\times10^{-4}$ | $9\times10^{-4}$ | *** |
| FR-ti-ta-ML vs. FR-ti-ta-HL | 15 | -0.22748 | $1.5\times10^{-4}$ | $1.35\times10^{-3}$ | ** |
| MR-ti-ta-FL vs. MR-ti-ta-ML | 14 | 0.50816 | $1.5\times10^{-4}$ | $1.35\times10^{-3}$ | ** |

| Comparison | N (flies) | Δ probability | $P_{raw}$ | $P_{adj}$ | Sig. |
|---|---|---|---|---|---|
| MR-ti-ta-FL vs. MR-ti-ta-HL | 14 | 0.07893 | 0.10165 | 1 | ns |
| MR-ti-ta-ML vs. MR-ti-ta-HL | 14 | -0.42923 | $1.5\times10^{-3}$ | 0.015 | * |
| HR-ti-ta-FL vs. HR-ti-ta-ML | 14 | 0 | 1 | 1 | ns |
| HR-ti-ta-FL vs. HR-ti-ta-HL | 14 | 0.66257 | $5\times10^{-5}$ | $4.5\times10^{-4}$ | *** |
| HR-ti-ta-ML vs. HR-ti-ta-HL | 14 | 0.66257 | $1\times10^{-4}$ | $9\times10^{-4}$ | *** |

**Figure 2H: First-SLC probability, across stimulation groups.** Two-sided unpaired permutation test on fly-wise first-SLC probability. Bonferroni family m = 9.

| Comparison | N (flies) | Δ probability | $P_{raw}$ | $P_{adj}$ | Sig. |
|---|---|---|---|---|---|
| FR-ti-ta-FL vs. MR-ti-ta-FL | 15/14 | -0.50523 | $5\times10^{-5}$ | $4.5\times10^{-4}$ | *** |
| FR-ti-ta-FL vs. HR-ti-ta-FL | 15/14 | -0.60843 | $5\times10^{-5}$ | $4.5\times10^{-4}$ | *** |
| MR-ti-ta-FL vs. HR-ti-ta-FL | 14/14 | -0.10321 | $1.5\times10^{-4}$ | $1.35\times10^{-3}$ | ** |
| FR-ti-ta-ML vs. MR-ti-ta-ML | 15/14 | 0.38388 | $2.5\times10^{-4}$ | $2.25\times10^{-3}$ | ** |
| FR-ti-ta-ML vs. HR-ti-ta-ML | 15/14 | -0.22748 | $5\times10^{-5}$ | $4.5\times10^{-4}$ | *** |
| MR-ti-ta-ML vs. HR-ti-ta-ML | 14/14 | -0.61136 | $5\times10^{-5}$ | $4.5\times10^{-4}$ | *** |
| FR-ti-ta-HL vs. MR-ti-ta-HL | 15/14 | 0.18213 | $5\times10^{-5}$ | $4.5\times10^{-4}$ | *** |
| FR-ti-ta-HL vs. HR-ti-ta-HL | 15/14 | 0.66257 | $5\times10^{-5}$ | $4.5\times10^{-4}$ | *** |
| MR-ti-ta-HL vs. HR-ti-ta-HL | 14/14 | 0.48044 | $5\times10^{-5}$ | $4.5\times10^{-4}$ | *** |

**Figure 2I: Fly-wise displacement direction.** Two-sided fly-wise directional displacement permutation test; statistic = angular distance between circular mean directions. Bonferroni family m = 9.

| Comparison | N (flies) | Δθ (°) | $P_{raw}$ | $P_{adj}$ | Sig. |
|---|---|---|---|---|---|
| FR-ti-ta-FL vs. MR-ti-ta-FL | 15/14 | 23.0728 | $1.5\times10^{-3}$ | 0.014 | * |
| FR-ti-ta-FL vs. HR-ti-ta-FL | 15/14 | 15.19139 | 0.012 | 0.108 | ns |
| MR-ti-ta-FL vs. HR-ti-ta-FL | 14/14 | 38.26419 | $5\times10^{-5}$ | $4.5\times10^{-4}$ | *** |
| FR-ti-ta-HL vs. MR-ti-ta-HL | 15/14 | 9.25348 | 0.035 | 0.315 | ns |
| FR-ti-ta-HL vs. HR-ti-ta-HL | 15/14 | 49.38494 | $5\times10^{-5}$ | $4.5\times10^{-4}$ | *** |
| MR-ti-ta-HL vs. HR-ti-ta-HL | 14/14 | 40.13146 | $5\times10^{-5}$ | $4.5\times10^{-4}$ | *** |
| FR-ti-ta-ML vs. MR-ti-ta-ML | 15/14 | 30.25847 | $5\times10^{-5}$ | $4.5\times10^{-4}$ | *** |
| FR-ti-ta-ML vs. HR-ti-ta-ML | 15/14 | 49.60353 | $5\times10^{-5}$ | $4.5\times10^{-4}$ | *** |
| MR-ti-ta-ML vs. HR-ti-ta-ML | 14/14 | 19.34506 | $5\times10^{-5}$ | $4.5\times10^{-4}$ | *** |

**Table S2. Comprehensive statistical comparisons and multiple-comparison adjustments related to Figure 5:** The statistical analysis is specified at the beginning of the block. Unless otherwise specified, the individual fly is the independent biological replicate and statistical unit. Across-group comparisons use two-sided unpaired permutation tests. All permutation tests use 20,000 permutations with a +1 correction. For each predefined comparison family, Bonferroni-adjusted P values were calculated as $P_{adj} = \min(P_{raw} \times m, 1)$, where m is shown in the block note. Significance notation: ns: $P_{adj} \geq 0.05$; *: $P_{adj} < 0.05$; **: $P_{adj} < 0.01$; ***: $P_{adj} < 0.001$.

**Figure 5B: Fly-wise landing latency (RMST).** Two-sided unpaired permutation test on fly-wise RMST (τ = 0.71 s). Bonferroni family m = 4.

| Comparison | N (flies) | Δ RMST | $P_{raw}$ | $P_{adj}$ | Sig. |
|---|---|---|---|---|---|
| Control vs. bristles | 15/7 | -0.61631 | $5 \times 10^{-5}$ | $4.5 \times 10^{-4}$ | *** |
| Control vs. all CS | 15/14 | -0.1291 | $6 \times 10^{-4}$ | $2.4 \times 10^{-3}$ | ** |
| Control vs. prx. CS2 | 15/18 | -0.04532 | 0.048 | 0.192 | ns |
| Control vs. prx. CS1 | 15/10 | -0.01017 | 0.119 | 0.476 | ns |
